# Characterization of hippocampal subfields using histology-based annotated postmortem MRI: Lessons for in vivo segmentation II

**DOI:** 10.1101/2025.11.21.689744

**Authors:** HL Tucker, B Karat, AT De Ruiter, D Parker, SA Lim, AE Denning, R Ittyerah, LM Levorse, W Trotman, A Bahena, E Chung, K Prabhakaran, ML Bedard, DT Ohm, L Xie, SR Das, JB Pluta, T Schuck, W Liu, S Pickup, E Artacho-Pérula, MM Iñiguez de Onzoño Martin, M Muñoz, FJM Romero, JC Delgado González, MM Arroyo Jiménez, M Marcos Rabal, AM Insausti Serrano, N Vilaseca González, S Cebada Sánchez, C de la Rosa Prieto, EB Lee, JA Detre, MD Tisdall, DJ Irwin, DA Wolk, DH Adler, R de Flores, D Berron, SL Ding, PA Yushkevich, CJ Hodgetts, LEM Wisse

## Abstract

High-resolution *in vivo* magnetic resonance imaging (MRI) of hippocampal subfields is a rapidly advancing field due to their implication in cognition, disorder, and disease. Hippocampal subfield segmentation on *in vivo* MRI is generally guided by postmortem reference material, which has been limited by small sample sizes that preclude comprehensive characterization of subfield border locations and their variability. Addressing this, we characterized hippocampal subfield border variability in two ultra-high-resolution postmortem MRI datasets with combined annotated histological sections, including cases with and without dementia. We examined: 1) the order of appearance and disappearance of subfields along the long axis of the hippocampus; 2) the order of appearance and disappearance of subicular subregions; 3) the medial-lateral position of subicular subregional boundaries along the hippocampal body; 4) the location of the CA3 relative to hippocampal head digitations; 5) the subfield borders in the hippocampal body relative to a volume proportion of the dark band; and 6) the association of hippocampal length and subiculum-CA1 border location with diagnosis, demographic factors, and factors related to postmortem imaging. Our findings reaffirmed that there is a consistent order of appearance and disappearance of subfields in the hippocampal head and tail, respectively. The subicular subregions exhibited a ‘first in, last out’ order of appearance and disappearance, and pre/parasubiculum consistently occupied half of the subicular complex in coronal slices throughout the hippocampal body. Hippocampal head digitations were not a reliable landmark for CA3 appearance, but SRLM proportionality did offer a potentially consistent approach for estimating CA2 and CA3 subfield borders in relation to the hippocampal border. No clear relationship was found between the anatomical features and diagnosis, demographic factors, and factors related to postmortem imaging. These findings have implications for the development and harmonization of hippocampal subfield segmentation protocols and interpretation of high-resolution functional MRI studies of the human hippocampus.

## 1. Introduction

Given the hippocampus’ key role in human memory and cognition (Moscovitch et al, 2016; Schachter et al, 2017, Zeidman & Maguire, 2016), and susceptibility to neurological and psychiatric conditions (Small et al, 2012; Thom, 2014; Harrison, 2004; Roeske, 2021; Busch et al, 2019), it has been a major neuroscience research priority to better understand its fine-grained structural and functional properties, particularly in relation to its subfields: namely, subicular complex, cornu ammonis (CA) 1-3, and dentate gyrus (DG) (Vos de Wael et al, 2018; Ding & van Hoesen, 2015; Insausti & Amaral, 2012; Moser & Moser, 1998). Animal studies, for example, have demonstrated differences in the functions, representations and computations supported by different hippocampal subfields, as well as variation in their vulnerability to aging and disease (Marr, 1971; Leutgeb et al, 2007; Knierim & Neunuebel, 2016; Ledergerber et al, 2021).

Translating these insights into the human brain has been facilitated by advancements in high-resolution MRI, which enable the study of hippocampal subfield structure and function in the living human brain (Zeineh et al, 2000; Zeineh et al, 2001). The subfields of the hippocampus are best delineated using high-resolution T2-weighted structural MRI (Yuskevich et al, 2015), which allows the visualization of the hypointense band – known as the stratum radiatum lacunosum moleculare (SRLM) – that separates the dentate gyrus from most of the CA subfields and the subicular complex (de Flores et al, 2019). Accurate registration of these high-resolution T2-weighted structural scans to high-resolution functional MRI data makes it possible to examine functional variation both within and across hippocampal subfield boundaries (Berron et al, 2016; Dimsdale-Zucker et al, 2018; Read, Berry et al, 2024), as well as to characterize broader structural and functional networks (Chang et al., 2021; Dalton et al, 2022). Structurally, high-resolution MRI has also enabled the detection of associations between subfield volumes and a range of cognitive functions, particularly in clinical populations. Such structural variations have been linked to symptom heterogeneity and may serve as markers of prodromal stages in neurological and psychiatric disorders (La Joie et al., 2013; Maruszak & Thuret, 2014; de Flores et al., 2015; Kerchner et al., 2010; Nuninga et al., 2020; Schobel et al., 2009; Twait et al., 2023).

However, the development of reliable, generalizable, and anatomically valid approaches to segmenting structural MR images is challenging, particularly as the boundaries between certain subfields (e.g., between CA1 and the subicular complex, and between CA3 and the dentate gyrus) are not easily distinguishable on in vivo MRI (Yushkevich et al, 2015). Typically, heuristics to guide segmentations and boundary placement have been obtained from anatomical atlases and reference material (Ding & van Hoesen, 2015; Mai et al, 2015; Amaral & Insausti, 1990; Insausti & Amaral, 2012; Duvernoy et al, 1998, 2005), but these often differ due to differences across neuroanatomical schools and/or inter-individual anatomical variability in the reference material itself (Wisse et al., 2017; Daugherty et al, 2025). Many anatomical atlases are based on a single or few cases, limiting their ability to capture population-level variation. The differences in hippocampal subfield border placement in these anatomical references likely contribute to divergent segmentation protocols for *in vivo* MRI and differing terminology (Yushkevich et al, 2015). Critically, such divergences may, in turn, account for discrepant results across studies (e.g., inconsistencies in how hippocampal subfields are affected by aging; see de Flores et al, 2015) and will necessarily affect the reproducibility of findings in both structural and functional MRI studies of hippocampal subfields.

In addition, anatomical atlases are mostly based on histological slices sectioned perpendicular to the anterior-posterior axis (but see Ding & van Hoesen, 2015, for an exception). This is different from T2-weighted MRI scans, which are typically acquired perpendicular to the hippocampal longitudinal axis with high in-plane but low out-of-plane resolution. The differences in orientation between histology and MRI can thus complicate the translation of histological information to neuroimaging contexts as both modalities are 2-dimensional, causing inaccuracies in the hippocampal subfield segmentation (de Flores et al, 2019; Supplementary Figure 1).

Thus far, these limitations have been partly addressed in a select few *ex vivo* MRI studies. One such study created an atlas of the hippocampal head using 15 postmortem brain cases, sliced perpendicularly to the hippocampal long axis (Ding & van Hoesen, 2015). Six of these medial temporal lobes, extracted from subjects with no clinical history and no presence of dementia pathology (ages 78-85), were selected as reference subjects. While this atlas showcased between-subject variability and images aligned with the orientation of MRI images, it did not include posterior regions of the hippocampus, nor did it evaluate potential hippocampal variability as a result of neurodegenerative conditions. In a later study conducted by de Flores et al (2019), nine brains obtained from donors with or without dementia were imaged with a 9.4 tesla (T) MRI and segmented based on co-registered annotated histological sections. The nine MRI scans, oriented to match the axis of in vivo MRI, were assessed to identify heuristics that could be translated to in vivo MRI segmentation protocols. Key findings were that (i) the sequence of subfield appearance was consistent in the hippocampal head; (ii) uniformity was observed in subfield composition in the posterior uncus but not the anterior uncus; (iii) the dark band visible on high-resolution T2 MRI corresponds only to CA-stratum lacunosum moleculare, and no other adjacent layers and strata; (iv) the border between the subicular complex and CA1 shifts to a more medial position along the anterior-posterior axis; and (v) the hippocampal tail resembles the body when resliced following its curvature.

These insights into the appearance and arrangement of histologically-defined hippocampal subfields are helpful for the interrogation of hippocampal subfields on in vivo structural and functional MRI, but key questions remain. First, as segmentations were acquired from a single neuroanatomical laboratory, it was not possible to examine possible disagreements in subfield border locations between neuroanatomy schools (Wuestefeld et al, 2024). Second, the study of de Flores et al (2019) treated the subicular complex - comprising of the subiculum (SUB), presubiculum (PrS) and parasubiculum (PaS) - as a single region, which is consistent with many in vivo segmentation schemes but obfuscates potential differences between the subregions (O’Mara et al, 2001). Finally, although some investigation into the role of demographic factors and diagnosis on the subiculum-CA1 border was made, further exploration could yield more nuanced information.

This paper aims to fill these gaps, expanding on the de Flores et al. (2019) paper by delving deeper into the original dataset as well as examining a more extensive dataset from 2024. The 2024 dataset features a broader spectrum of neurodegenerative disease diagnoses, including additional detail on neuropathological diagnosis, allowing for greater generalizability. Furthermore, the 2024 dataset was segmented to include labels for subicular complex regions, i.e. the subiculum, presubiculum, and parasubiculum. Expanding on these subregions and examining their properties in relation to hippocampal macroanatomy offers the potential to develop more refined segmentation schemes and thus more detailed examination of subicular complex structure and function in humans (Dalton et al, 2017; Read, Berry et al, 2024). Finally, the inclusion of a second, more comprehensive dataset allows for comparisons of subfield border locations across laboratories. Thus, this paper aimed to provide an analysis of the location of hippocampal subfield borders in relation to anatomical landmarks, as well as demographic, diagnostic and postmortem imaging factors, with the overarching aim of informing in vivo segmentation approaches. Specifically, the following topics are addressed:

- The order of appearance of hippocampal subfields along the long axis of the hippocampal head
- The order of disappearance of hippocampal subfields along the long axis of the hippocampal tail
- The order of appearance and disappearance of subicular complex subregions along the long axis of the hippocampus
- The boundary locations of subicular complex subregions in coronal slices in the hippocampal body
- The location of CA3 relative to hippocampal head digitations
- The borders of subfields in coronal slices in the hippocampal body relative to a volume proportion of the SRLM around the DG
- Hippocampal length as a function of demographic factors and factors specific to postmortem imaging
- Relative location of SUB-CA1 border in coronal slices in the hippocampal body as a function of demographic factors and factors specific to postmortem studies

## 2. Methods

### 2.1. Donor cohorts

All cases were obtained in accordance with local laws and ethical review authorities. Postmortem consent from next-of-kin was always obtained, and antemortem consent from the patient was obtained where possible. Overall, the datasets consisted of 26 cases, of which 9 were women and 17 were men, where the mean age at death was 75.5 years (± 9.2 years). The cases were comprised of 12 left hemispheres and 14 right hemispheres.

*2019 dataset:* The 2019 dataset contained nine samples, two from autopsies performed at the University of Pennsylvania Center for Neurodegenerative Disease (CNDR) and seven from the National Disease Research Interchange (NDRI) brain bank, with ages ranging from 60-90 years (de Flores et al, 2019; Adler et al, 2018). See Table 1 for demographics.

**Table 1.** Demographics of the 2019 dataset (N = 9)

| Sample ID | Hemisphere | Age (years) | Sex | Clinical Diagnosis | Fixation (days)<sup>d</sup> | PMI (hours) |
| --- | --- | --- | --- | --- | --- | --- |
| NDRI01<sup>a</sup> | R | 89 | F | OD | 108 | 9.5 |
| NDRI02 | R | 76 | M | NC | 256 | 24 |
| NDRI05 | L | 61 | M | AD | 66 | 21 |
| NDRI08 | R | 75 | M | NC | 523 | 15 |
| NDRI09 | R | 78 | F | NC | 217 | 18 |
| NDRI12 | L | 81 | F | NC | 527 | 11.4 |
| NDRI14 | L | 90 | F | AD | 442 | 13 |
| CNDR02<sup>b</sup> | R | 60 | M | AD | 82 | 12 |
| CNDR10<sup>c</sup> | L | 73 | M | OD | 22 | 19 |

<sup>a</sup>All measurements were performed in 0.2x0.2x0.2 mm<sup>3</sup> but the acquired resolution was 0.16x0.16x0.16 mm<sup>3</sup>. <sup>b</sup>Pathological diagnosis: High Alzheimer's disease neuropathologic change. B score is 3. <sup>c</sup>Pathological diagnosis: Progressive supranuclear palsy, low Alzheimer's disease neuropathologic change. B score is 1. <sup>d</sup>Fixation time at time of MRI. AD=Alzheimer's disease; NC=Non-dementia control; OD=Other dementia

*2024 dataset:* The 2024 dataset consisted of 17 samples, 5 obtained from the University of Pennsylvania CNDR (non-overlapping with the two from the 2019 dataset) and 12 from the Human Neuroanatomy Laboratory (HNL) brain bank at the University of Castilla-La Mancha, with ages ranging from 45-93 years (Ravikumar et al, 2021). The CNDR samples in the 2024 dataset were obtained from dementia research patient donors while the University of Castilla-La Mancha samples were mostly older adults with no known neurological diseases from the surrounding geographical area. See Table 2 for the demographics.

**Table 2.**
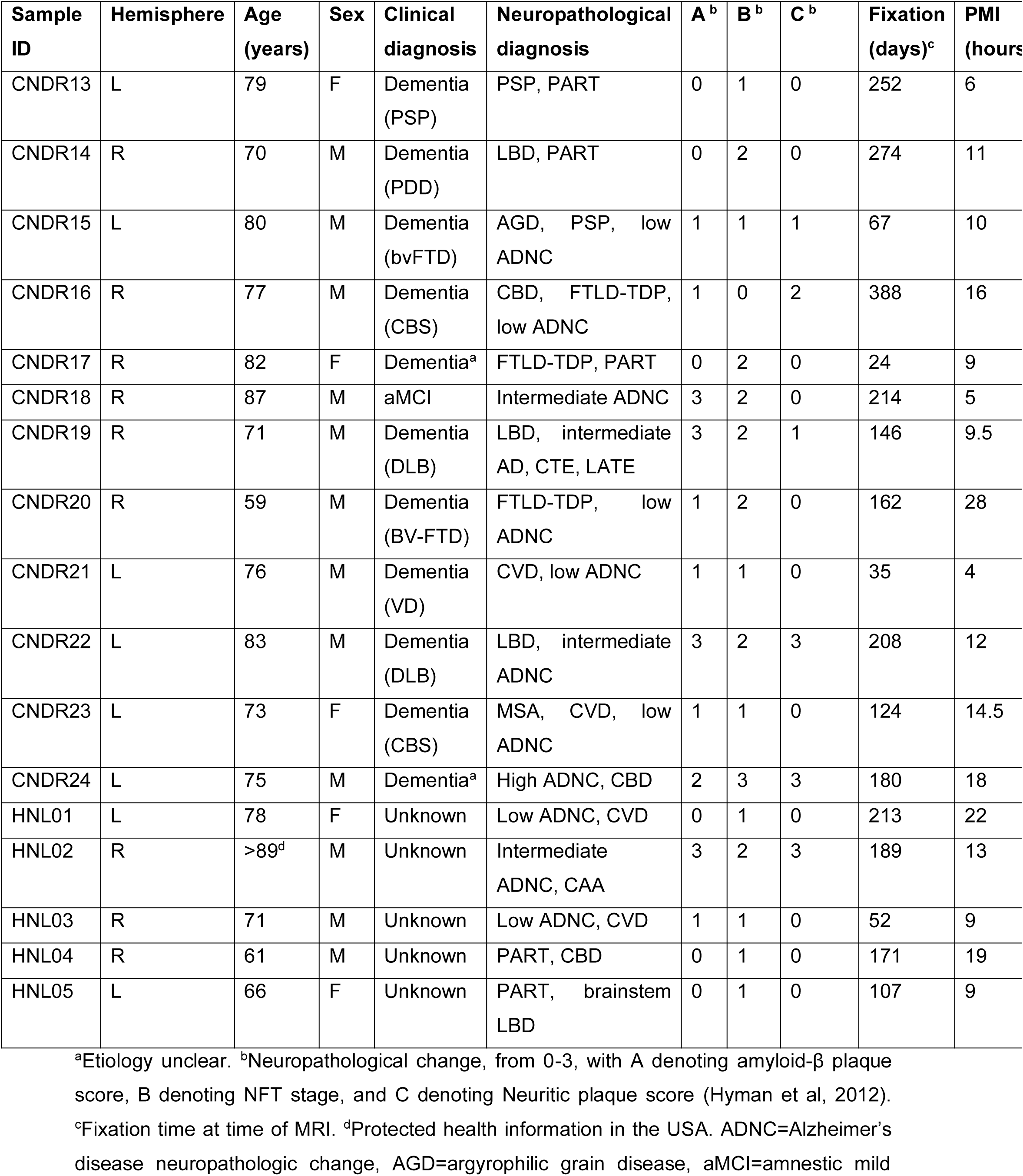
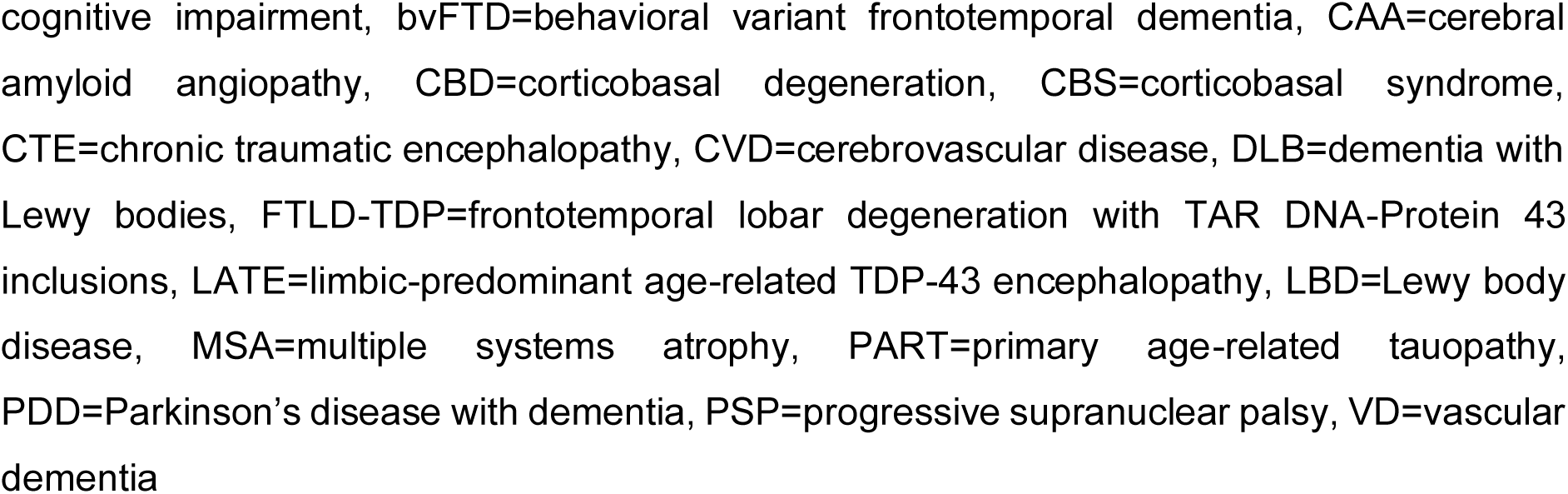
Demographics of the 2024 dataset (N = 17)

### 2.2. Imaging procedures

In both datasets, tissue from one hemisphere was sampled for imaging and the other for histology and neuropathological diagnosis. At CNDR, the examined hemisphere was alternated every autopsy. The hemispheres at HNL were also selected at random for imaging (Yushkevich et al, 2021). The hemispheres selected for postmortem imaging were fixed for a minimum of 21 days. Cases were then imaged using a Varian 9.4T 31-sm bore scanner at 0.2 x 0.2 x 0.2 mm^3^ resolution before histological processing.

*2019 dataset:* The cases were embedded in paraffin and sectioned into 5 µm thick sections with 200 µm spacing and stained with the Klüver-Barrera method (Klüver & Barrera, 1953). The final sections were digitally scanned at 0.5 x 0.5 µm resolution and aligned with the MRI images using the in-house developed software HistoloZee (Adler et al., 2018).

*2024 dataset:* Cases from the 2024 dataset were also imaged in a 7T human MRI scanner at 0.4 x 0.4 x 0.4 mm^3^ resolution as an addition to the 9.4T MRI scan. For dense serial histology, a custom sectioning mold was 3D printed using the 7T MRI scans. This mold guided tissue sectioning orthogonally to the main axis, which in turn facilitated registration between histology and MRI, yielding circa four 20 mm thick blocks which were then frozen. Block-face images were captured using an overhead camera. Blocks were then sectioned into 50 µm sections using a sliding microtome, with no gaps between sections. Every 10^th^ 50 µm section was thionin stained for the Nissl series. The Nissl series contained circa 40 sections per block at 0.5 mm intervals. All sections were mounted on 75 mm x 50 mm glass slides, which were then scanned with a pixel size of 0.4 µm at 20x resolution. These digital scans were uploaded to an internal, digital histology archive that aids with web-based visualization and labelling of anatomical structures.

### 2.3. Segmentation of hippocampal subfields

*2019 dataset:* Hippocampal subfield boundaries were annotated in each histological section using Ding and van Hoesen’s (2015) anatomical protocol and later reviewed by an expert neuroanatomist (SLD) for validation. The annotated histology sections were then registered to the 9.4T postmortem MRI using a graph-theoretic automated histology reconstruction algorithm, described by Adler et al (2014), and utilized to guide the segmentations of hippocampal subfields in MRI space using ITK-SNAP (Yushkevich et al, 2006). The DG, CA1, CA2, CA3, and SUB were segmented through this process. The exception to this segmentation approach was the dark band, which was specifically segmented based on the features visible on MRI using a semi-automated approach (Adler et al, 2014).

*2024 dataset:* The hippocampal subfields (DG, CA1-3, SUB, PrS, PaS) were annotated on histological sections by expert neuroanatomists following the criteria established by Insausti and Amaral (2012; Insausti et al, 2017; Rosene & Van Hoesen, 1987), in an open-sourced web-based system using images of Nissl slides. The histological sections were then registered to the 9.4T postmortem MRI using a multi-step largely automated pipeline, described in more detail in Yushkevich et al (2021). The annotations were superimposed on the co-registered MRI and histology images to guide manual segmentation of hippocampal subfields in MRI space in ITK-SNAP. Again, the dark band was segmented based on features visible on MRI using a semi-automated approach (Yushkevich et al, 2021).

Note that in both datasets the segmentations do not cover the full length of the hippocampus on postmortem MRI. This was the result of missing histology data, gaps between blocks due to shrinkage of the postmortem tissue, a slightly different angulation of the co-registered histology sections therefore not covering the full coronal MRI slice and tears and other artefacts on the histology sections. As the segmentations on the postmortem MRI were copied one-on-one from the histology sections, missing histological data led to missing portions either in-plane or on several slices on the postmortem MRI. This led to exclusion of some cases for the separate research questions, specified in each of the sections and Supplementary Table 1.

### 2.4. Analyses

As this paper covers a breadth of research questions which require different approaches, the method for each analysis is noted within each subsection of the results. Which cases are used for each question is shown in Supplementary Table 1. Unless stated differently, measurements were completed in ITK-SNAP (Yushkevich et al, 2006).

## 3. Results

The following results section details the individual aims, approaches and results of each research question, along with a brief discussion of the implications of each finding. A summary of the main findings per subsection can be seen in Table 3.

**Table 3.** Summary table of the main findings per results-section.

| Section | Aim | Main finding |
| --- | --- | --- |
| 3.1 | To provide a detailed description of the order of appearance of subfields within the hippocampal head. | Subfields appeared in the following order in most, but not all, cases: SUB-CA1-DG-CA3-CA2. There is variability in the order of CA3 and CA2. |
| 3.2 | To characterize the order of disappearance of subfields within the hippocampal tail. | The SUB and CA1 are present in the most posterior 2 mm or 1 mm of the hippocampus, but there is more variability in other subfields. |
| 3.3 | To characterize the emergence and disappearance of the subicular complex regions. | Subicular complex regions consistently appear in the following order: SUB-PrS-PaS. PrS consistently emerges closely after CA1. Subicular complex region disappear in a 'last in, first out' order. |
| 3.4 | To investigate the location of the PrS and PaS borders in relation to the length of the subicular complex throughout the hippocampal body. | The PrS-SUB border is consistently located halfway through the subicular complex. The PaS occupies the smallest proportion of the subicular complex, followed by the PrS, with the SUB occupying the largest proportion. |
| 3.5 | To characterize the location of CA3 relative to digitations in the hippocampal head. | CA3 tends to appear in-between digitations, but the exact location varies considerably. |
| 3.6 | To investigate if subfields project onto a consistent relative volume proportion of the SRLM in a way which is useful for in vivo segmentation. | The SRLM proportionality measure was a consistent tool for quantifying the relative size of the CA2 and CA3, but more variable for CA1. |
| 3.7 | To investigate if demographic factors and factors specific to postmortem imaging are associated with different measures of hippocampal length. | There were no significant associations between hippocampal length measures and demographic factors or factors specific to postmortem imaging. |
| 3.8 | To investigate the SUB-CA1 border location in the hippocampal body and if this position is associated with demographic factors and factors specific to postmortem imaging. | The SUB-CA1 border location in the hippocampal body shifts medially along the anterior-posterior axis and is more variable in the posterior part of the hippocampal body. There were no significant associations of CA1/SUB border location and demographic variables or factors specific to postmortem imaging. |

SUB = subiculum, CA = cornu ammonis, DG = dentate gyrus, PrS = presubiculum, PaS = parasubiculum, SRLM = stratum radiatum lacunosum moleculare.

### 3.1. The order of appearance of hippocampal subfields along the long axis of the hippocampal head

*Aim:* The aim of this section was to provide a detailed description of the order in which subfields appear within the hippocampal head, replicating the approach of de Flores et al. (2019) within the larger 2024 atlas. Furthermore, we evaluated whether the order of appearance differs between the 2024 and 2019 atlases, providing an opportunity to examine between-lab variations in border definitions. This section focused on the CA subfields, the DG, and the SUB. The appearance of the PrS and PaS is examined separately in Section 3.3.

*Approach:* This analysis included 14 out of 17 specimens from the 2024 atlas and three out of nine from the 2019 atlas. Specimens were excluded if there was uncertainty regarding the exact starting point of a subfield. This arose when the annotations anterior to the first labelled slice of a subfield contained missing sections. It was possible, therefore, that the subfield appeared more anteriorly (Supplementary Figure 2). To address this, we excluded cases if there were five or more consecutive slices with insufficient coverage anterior to the appearance of any subfield.

The appearance of the SUB was defined as the slice containing the anterior tip of the hippocampus as visualized on MRI, independent of segmentation. The start of each CA subfield was determined by the first coronal slice showing four contiguous voxels labelled as CA1, with contiguous voxels connected by either corners or edges. In cases where the appearance of a CA field was preceded by slices with incomplete coverage, the four-voxel rule was not applied. Instead, a single segmented voxel was considered sufficient to indicate the appearance of the subfield. The DG, identifiable solely by MRI features, was defined as the first slice where grey matter appeared fully surrounded by the dark band, even if no segmentation label was visible or had been present in previous slices. Consistent with de Flores et al. (2019), the uncal apex was used to define the end of the hippocampal head, which marked the most posterior slice where any uncal tissue remained visible (Poppenk et al., 2013). The distances between regions were calculated by multiplying the number of slices by the slice thickness (0.2 mm) and converting them into percentages of the hippocampal head’s total length, defined by the distance from the anterior tip of the hippocampus to the posterior tip of the uncal apex. See the Supplementary Methods section for some additional notes on segmentation inconsistencies.

*Results:* In the 2024 atlas, 11 out of 14 cases showed the appearance order SUB-CA1-DG-CA3-CA2 (Figure 1; Figure 2 for the order plotted per case). In three cases, CA3 appeared before the DG, while the order of the other subfields remained consistent with the typical pattern. For the ‘canonical’ order, the average distances between subfields relative to the preceding subfield across all cases were as follows (Supplementary Table 2): CA1 appeared 1.9 mm (11.3% of the head length) posterior to the SUB, the DG appeared after 3.7 mm (22.1%), CA3 after 1.6 mm (9.5%), and CA2 after 3.0 mm (18.8%). In the three cases where CA3 appeared before the DG, CA3 emerged 3.3 mm (18.6%) after CA1, and the DG followed after 0.8 mm (4.7%).

**Figure 1.**
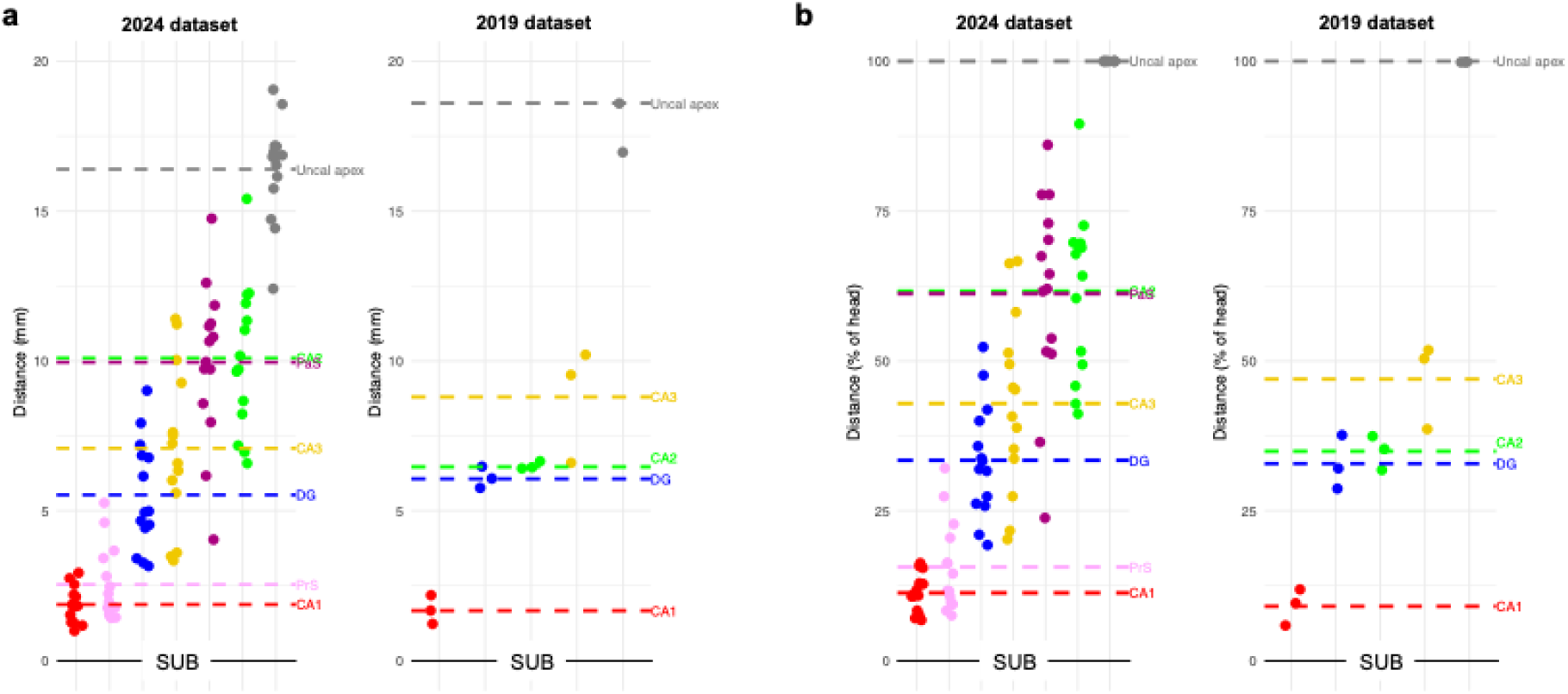
The order of appearance of hippocampal subfields (and subicular complex regions; see Section 3.1 & 3.3) is shown in a) millimeters (from most anterior hippocampal slice), and b) percentages (of head length). Zero on the y-axis corresponds to the most anterior slice (labeled as subiculum, SUB) of the hippocampus. Each data point relates to a specific case. The dashed lines indicate the averages for the group. Order of appearance for individual cases is presented in Figure 2. SUB = subiculum, CA =cornu ammonis, DG = dentate gyrus, PrS = presubiculum, PaS = parasubiculum.

**Figure 2.**
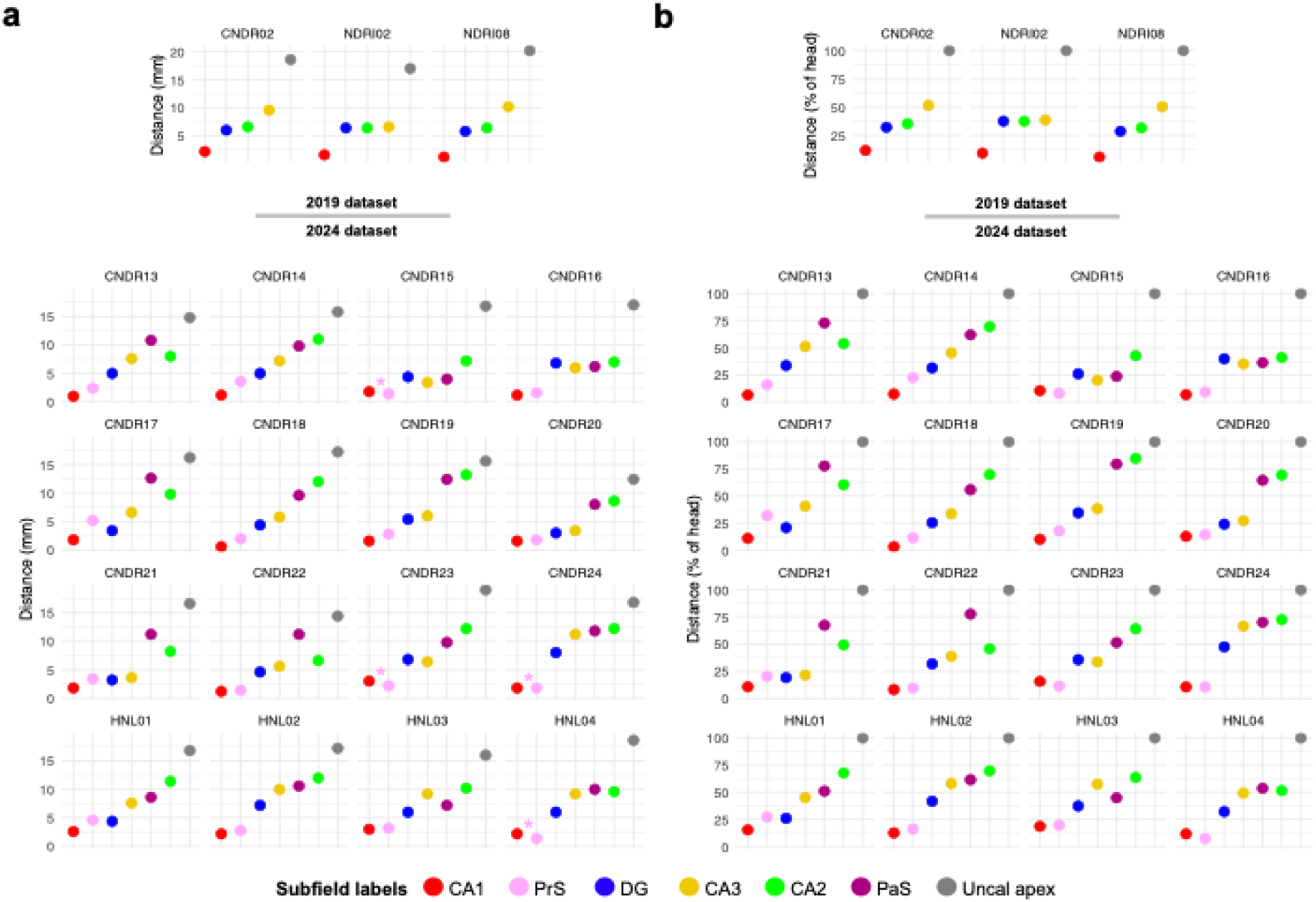
The order of appearance of hippocampal subfields in the head in each case of the 2024 dataset. All subfields are shown in the ‘canonical’ order for this sample, namely SUB (point zero on the y-axis), CA1, PrS, DG, CA3, CA2, PaS, and uncal apex. SUB = subiculum, CA = cornu ammonis, DG = dentate gyrus, PrS = presubiculum, PaS = parasubiculum.

For the 2019 atlas, two out of three cases demonstrated the appearance order SUB-CA1-DG-CA2-CA3, consistent with the findings of de Flores et al. (2019). In the third case, the DG and CA2 emerged simultaneously on the same coronal slice, whereas the remaining subfields followed the standard pattern. For the standard order, the average distances between subfields across all cases were as follows: CA1 appeared after 1.7 mm (9.1% of the head length) following the SUB, the DG appeared after 4.4 mm (23.8%), CA2 after 0.4 mm (2.1%), and CA3 after 2.3 mm (12.0%). Compared to the 2024 atlas, the order of CA2 and CA3 appearance was reversed.

*Discussion:* Here, we provide a detailed description of the order in which subfields appear along the long axis of the hippocampus. For the 2024 atlas, the analysis shows that in the majority of the cases, the subfield appearance order was SUB-CA1-DG-CA3-CA2. In three specimens, CA3 emerged before the DG, whereas the order of the other subfields remained consistent with the predominant pattern. The 2019 atlas revealed a similar subfield order, also reported by de Flores et al. (2019), with the notable exception that CA3 appeared after CA2 in all cases.

The findings suggest both inter-individual consistency for most of the subfields and cases but also some variability in subfield appearance order. For example, some cases in the 2024 atlas showed a reversal where the DG appeared (0.8 mm) posterior to CA3, contrary to the typical order. This reversal may reflect individual differences in hippocampal anatomy and could be relevant for in vivo segmentation, depending on the slice thickness used. Further research with a larger sample size could help determine whether the alternative order has biological significance, especially because it contrasts with established neuroanatomical references, such as those presented by Mai et al. (2015), which typically describe the DG as preceding CA3.

Notably, in the 2024 atlas, CA3 consistently appeared anterior to CA2, while in the 2019 atlas, the reverse was observed, suggesting a discrepancy between the two atlases. A likely explanation is that the atlases were annotated/guided by neuroanatomists from different schools, each of which may emphasize distinct features when defining the cytoarchitectonic fields of the hippocampus. A recent study by Wuestefeld et al. (2024) found lower levels of agreement among neuroanatomists from different schools when annotating transitional zones within the medial temporal lobe. Although their study focused on the medial temporal lobe cortex, similar challenges may arise in hippocampal delineation in transitional zones such as the anterior border of CA2 and CA3. Transitions between subfields are often gradual rather than discrete, and the criteria for delineation may vary depending on which features are prioritized. While the small sample size limits definitive conclusions about consistent differences in CA2 and CA3 order between atlases – and, thus, between neuroanatomical schools – previous cytoarchitectonic studies have shown variability in the relative positioning of these fields, recapitulating our findings (CA2 anterior to CA3, Ding & Van Hoesen, 2015; CA3 anterior to CA2, Insausti & Amaral, 2012).

In general, this section offers valuable insights into the in vivo segmentation of the hippocampal head. Specifically, the SUB is the most anterior subfield followed by CA1 in all cases, followed by the DG in most cases. Meanwhile, the order of CA2 and CA3 is less clear. However, the error introduced by any heuristic regarding the order of CA2 and CA3 will be relatively small because the distance between CA2 and CA3 was on average between 2.3 and 3.0 mm (equivalent to 1.5 x 2 mm slices). Although minor discrepancies in segmentation may not significantly impact broader research on hippocampal subfields, they may be critical for detailed anatomical studies of the hippocampal head and clinical applications requiring high precision. A consensus between neuroanatomy schools may be useful to resolve this discrepancy in CA2-CA3 order.

### 3.2. The order of disappearance of hippocampal subfields along the long axis of the hippocampal tail

*Aim:* The aim of this section was to characterize the order in which subfields disappear along the long axis of the hippocampal tail. This aim replicates the study by de Flores et al. (2019) with the larger 2024 atlas. Additionally, we assessed whether the order of disappearance differs between the 2024 and 2019 atlases.

*Approach:* Considering the limited number of cases with histological sections covering the entire hippocampal tail, we adopted the methodology used by de Flores et al. (2019). Specifically, we identified the most posterior slice of the hippocampus (Supplementary Figure 3) and described the subfields both 2 mm and 1 mm anterior to this slice to match common slice thicknesses on *in vivo* MRI. The analysis included 10 out of 17 cases from the 2024 atlas and six out of nine from the 2019 atlas, excluding seven and three cases, respectively. Furthermore, five additional cases from the 2024 atlas and two from the 2019 atlas were excluded from the analysis focusing on the most posterior 1 mm of the tail. Cases were excluded if the most posterior slice could not be determined or estimated with reasonable confidence or if the slice to be analyzed was (partially) cut off or contained insufficient segmentation coverage.

Unlike the approach employed in Section 3.1, we primarily relied on histology-based segmentations rather than MRI features to identify subfields in the posterior hippocampus, with a few exceptions. See the Supplementary Methods for section 3.2 for additional details.

*Results:* In the 2024 atlas, at 2 mm anterior to the most posterior slice, the subicular complex and CA1 were present in 100% of cases, CA3 in 70%, the DG in 60%, and CA2 in 50% (Figure 3a). Shifting 1 mm posteriorly, the SUB and CA1 were identified in 100% of cases, CA3 in 60%, CA2 in 40%, and the DG in 20%. In the 2019 atlas, at 2 mm anterior to the most posterior slice, the SUB, CA1, the DG, and CA2 were present in 100% of cases, while CA3 was identified in 83.3% (Figure 3b). Moving 1 mm posteriorly, CA1 was present in 100% of cases, with the subicular complex, CA2, and CA3 each identified in 75% and the DG in 50%. The results align with the findings of de Flores et al. (2019).

**Figure 3.**
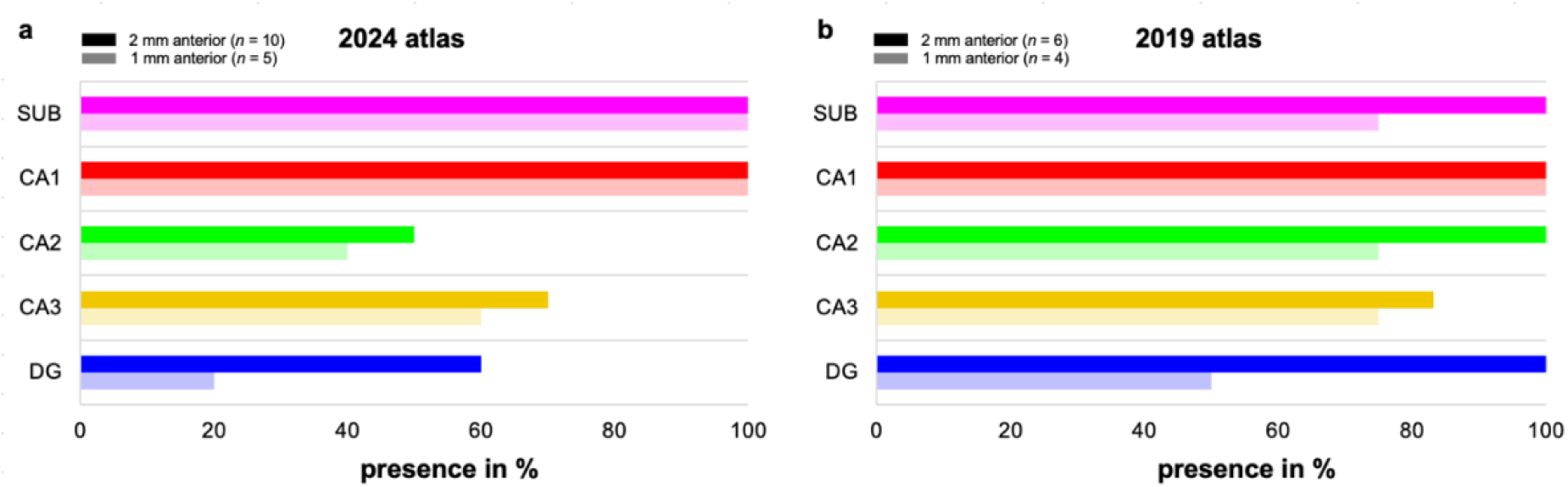
The distribution of subfield presence in the hippocampal tail in the (a) 2024 and (b) 2019 atlases. The figure shows the percentages of subfields (the SUB, CA1, CA2, CA3, and the DG) present at both 2 mm and 1 mm anterior to the most posterior slice. SUB = subiculum, CA = cornu ammonis, DG = dentate gyrus.

*Discussion:* For the characterization 2 mm typical slice thickness for high-resolution T2-weighted 3T MRI scans, anterior to the most posterior tail slice, the analysis showed that the SUB and CA1 were present in 100% of cases, while CA3 was identified in only a subset of cases in both atlases. Results for CA2 and the DG were more variable, with these subfields present in 100% of cases in the 2019 atlas but only in about 50%–60% of cases in the 2024 atlas. For the characterization 1 mm anterior to the most posterior tail slice (approximately the slice thickness in high-resolution T2-weighted 7T MRI scans), CA1 was present in all cases, and the SUB was observed in almost all cases (75%–100%) in both atlases. In contrast, the presence of the DG, CA2, and CA3 was more variable and observed in a smaller portion of cases. Interestingly, CA2, CA3, and the DG occurred more often in the posterior 1 mm of the hippocampal tail in the 2019 atlas compared to the 2024 atlas.

The findings suggest that the subicular complex and CA1 can be reliably segmented until the end of the hippocampus using slices that are either 2 mm or 1 mm thick. However, the variability observed in the presence of CA3, the DG, and CA2 in the hippocampal tail may have implications when segmenting these subfields across the entirety of the hippocampal tail. The DG benefits from a distinct macro-anatomical landmark—the SRLM or dark band—which should ideally be used to guide its segmentation. Even with 2 mm slices, due to the observed variability, continuing to segment the DG until the SRLM is no longer traceable may help to ensure accuracy. The inconsistent appearance of CA2 and CA3 at both 2 mm and 1 mm anterior to the most posterior slice suggests that reliably segmenting these subfields up to the more posterior hippocampal boundary may not be feasible, irrespective of slice thickness.

The difference in the presence of CA3, the DG, and CA2 between the two atlases at both the posterior 2 mm and 1 mm of the hippocampal tail may be attributed to variations in inherent neuroanatomy. Additionally, differences in annotation practices between atlases—and, by extension, between neuroanatomical schools—may also have contributed to this variability. As noted by de Flores et al. (2019), the composition of subfields in the posterior hippocampus visible on MRI can be influenced by the tail’s curvature, and re-slicing of the tail could potentially reduce some of the observed discrepancies.

### 3.3. The order of appearance and disappearance of subicular complex subregions in the hippocampus

*Aim:* The majority of hippocampal segmentation protocols for in vivo MRI (at both 3T and 7T) have adopted an inclusive ‘subiculum’ region-of-interest, which extends typically from the distal border of CA1 to the superior border of the entorhinal and parahippocampal cortices (Yushkevich et al., 2015; but see Dalton et al., 2017). This broader ‘subiculum’ region, more appropriately labeled ‘subicular complex’ (see O’Mara et al., 2001), encompasses several cytoarchitectonically defined subregions, including subiculum proper (SUB), presubiculum (PrS), and parasubiculum (PaS)^1^ (Ding, 2013).

Importantly, differentiating these subicular complex regions has potential relevance to understanding human memory and spatial cognition. For instance, electrophysiological studies in rodents have identified specialized cell types across different regions of the subicular complex, including head direction cells in the PrS/PaS (Ranck, 1994; Taube, 1990), place cells in the SUB (Sharp, 2006), and border cells throughout the subicular complex (Lever et al., 2009). Human fMRI data also suggest that a region likely corresponding to anterior PrS/PaS supports visual scene perception and memory (e.g., Dalton & Maguire, 2017; Grande et al., 2023; Hodgetts et al., 2017; Read et al., 2024).

In this section, we aimed to characterize the emergence of subicular complex regions, i.e. the PrS and PaS within the hippocampal head, as well as their appearance relative to other hippocampal subfields, including the SUB (outlined in detail in Section 3.1). Further, we explored their order of disappearance with respect to the most posterior point of the hippocampus (see Supplementary Methods Section 3.3). Such observations may be relevant to guiding future segmentation protocols for in vivo MRI (i.e., by guiding the development of new heuristics), and will provide important information relevant to interpreting structural and functional MRI findings within the subicular complex.

*Approach:* As in Section 3.1, 14/17 cases in the 2024 dataset were used to characterize the appearance of subicular complex subregions within the hippocampal head. The most anterior and posterior slice of the hippocampus were defined as described in Sections 3.1 and 3.2 respectively. The appearance of a given subicular complex substructure was defined as any slice containing 4 or more contiguous voxels for that substructure (connected by either edges or corners). The most posterior slice in which the PrS and PaS were segmented was noted for each case and its distance to the tail endpoint was calculated – both in mm and as a proportion of whole hippocampal length.

For the disappearance analysis, one additional case (HNL01) was removed as the most posterior slice could not be determined. The three additional cases excluded in section 3.2 (HNL02, HNL04 & HNL05) were retained in this section, as the lack of segmentation coverage in the hippocampal tail did not affect the slices containing PrS and PaS. This resulted in a final sample of 13 cases for the disappearance analysis.

*Results:* The order of appearance of subicular complex regions – relative to each other – was consistent across all subjects (Figure 1): subiculum constituted the most anterior slice of the hippocampus (Section 3.1), followed by the PrS 2.5 mm posterior to this (15.6% of the head length). The PaS then appeared 7.4 mm posterior to the PrS (45.6%). The PaS appearance point also corresponded to 10 mm posterior to the most anterior slice of the hippocampus - or 61.2% of total head length (see Figure 1; Figure 2 for order of appearance plotted by case).

While the emergence of subicular complex regions was consistent across cases (SUB > PrS > PaS), there was greater inter-individual variability with respect to their ‘nearest neighbors’ (Figure 2). Notably, PrS and CA1 were found to emerge at similar points along the hippocampal head across cases (Figure 2). On average, the PrS was found to emerge only 0.67 mm posterior to the appearance of CA1 (4.3%). However, this was not seen in all participants, with N = 4 samples possessing PrS slices *anterior to* CA1 (shown by pink asterisks in Figure 2). The distance between PrS and DG was considerably larger than that seen with CA1, and also more variable across cases. On average, DG was found to appear 3 mm *posterior to* the emergence of PrS, corresponding to 17.8% of head length. Despite this average pattern, 3 samples still showed DG slices *anterior to* PrS.

Relative to PaS, CA3 appeared anteriorly in all samples with an average distance of 2.8 mm (18% of head length). PaS and CA2 were found to appear at a similar point on average, with CA2 appearing only 0.14 mm posterior to PaS (0.42%). Five cases did, however, have PaS appearing posterior to CA2; these flipped cases also tended to have larger distances, with an average PaS-to-CA2 distance being 2.28 mm (14.7%) in these cases versus 1.5 mm (8.8%) in the canonical group.

In terms of disappearance, the subicular complex regions displayed a ‘last in, first out’ pattern (see Figure 4). The PaS, which appeared most posteriorly in the hippocampal head, disappeared 16.2 mm anterior to the hippocampal tail endpoint, a distance that corresponds to 7.6% of total hippocampal length. As a result of it more posterior point of appearance in the head (see above), and its relatively anterior disappearance, the PaS extended only 16.4 mm in length, which corresponds to 37.9% of total hippocampal length.

**Figure 4.**
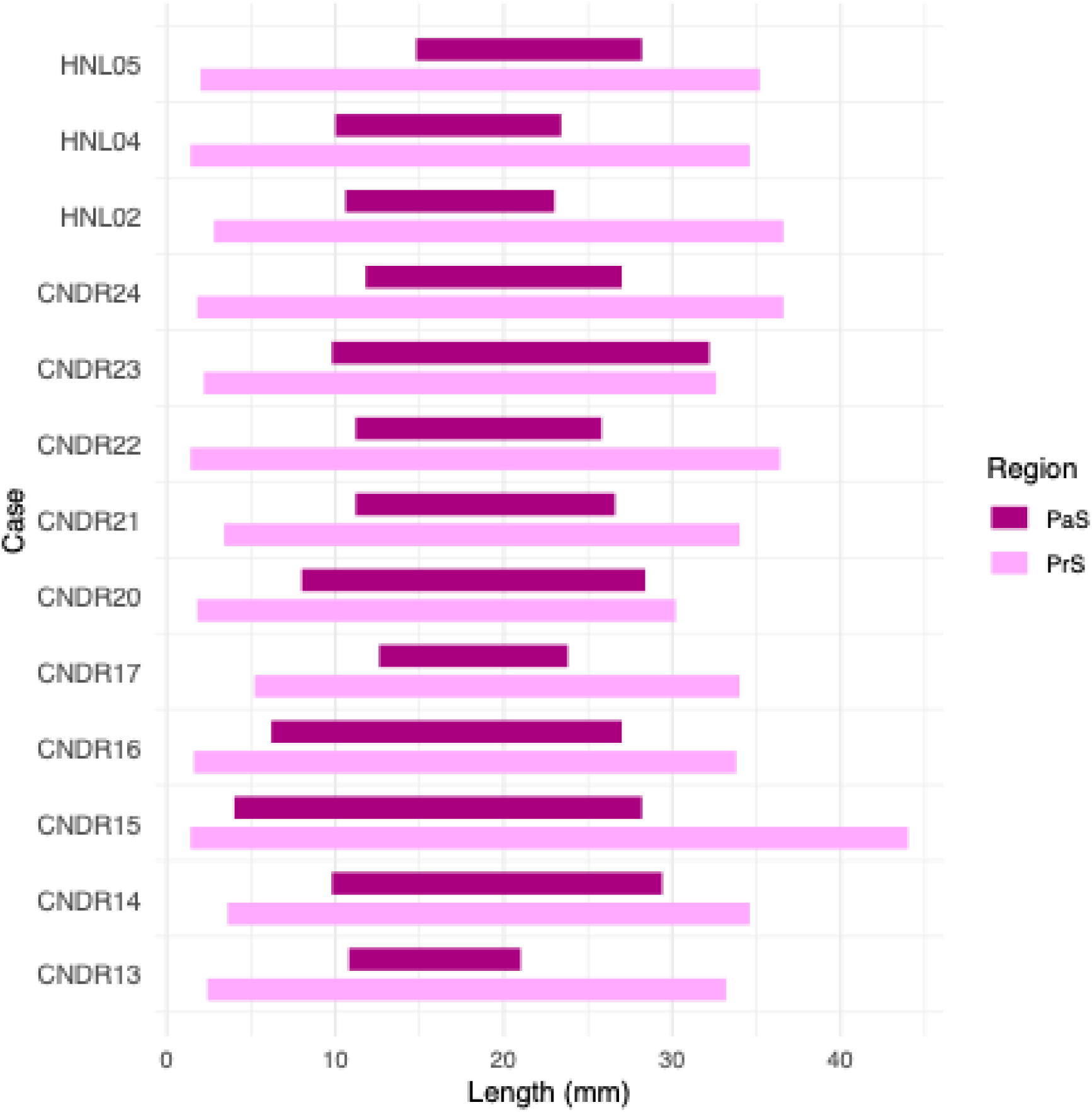
The length of the PaS and PrS within the hippocampal body, showing the points of appearance and disappearance of the subicular complex regions along the longitudinal axis. PaS = Parasubiculum, PrS = Presubiculum.

The PrS disappeared more posteriorly in all cases, with segmentation ending on average 8.6 mm posterior to PaS, and 7.6 mm anterior to the tail endpoint (17.8% of hippocampal length). In only one case (CNDR15) did the PrS extend to 2mm anterior to the most posterior hippocampal slice. Given its appearance near the most anterior point of the hippocampus (2.5mm), the PrS spanned much of the hippocampus, with a mean length of 32.7 mm, equivalent to 76.6% of total hippocampal length.

*Discussion:* These analyses, conducted within 14 cases, revealed that the subicular complex regions appeared in a consistent order across all samples: namely, SUB was visible in the most anterior slice of the hippocampal formation, followed by the PrS, and then the PaS posterior to this. Notably, the anterior appearances of the SUB and PrS were relatively close together, whereas the PaS typically appeared beyond the halfway point of the hippocampal head. In all but two of the cases, the PrS was present by the 4mm mark, suggesting that potential segmentation schemes for subicular complex regions could contain the PrS – with reasonable confidence – from the third 2 mm slice onwards.

In relation to non-subicular regions, the PrS was found to emerge less than 1 mm posterior to CA1, and this CA1-PrS ordering was fairly consistent across cases. On this basis, in vivo segmentation rules for the emergence of CA1 could be feasibly extended to the PrS – particularly given the relatively low inter-sample variability in where these regions appear (e.g., “begin segmenting PrS on the slice immediately posterior to the emergence of CA1”). In addition, the majority of cases (11/14) showed PrS slices located anterior to DG, confirming a relatively canonical anterior-to-posterior pattern of SUB-CA1-PrS-DG in the head of the hippocampus.

The PaS was located posterior to CA3 in all cases examined, and was typically just anterior to CA2. There was, however, large variability across cases, and in five cases the reverse pattern was observed (CA2 *anterior to* PaS), which also corresponded to larger distances between them. For MRI images with a 1-2 mm out-of-plane resolution, a reasonable recommendation would be to segment PaS at the point where CA2/3 are segmented in the hippocampal head. However, it is unclear how PaS relates to CA2 and CA3 in segmentations where the order of CA2 and CA3 is reversed (see Section 3.1).

Finally, we observed that subicular complex regions followed a ‘last in, first out’ pattern, with the PaS disappearing first, followed by the PrS, and then finally SUB. Given its late appearance, the PaS was segmented across relatively few slices overall (16.6 mm), corresponding to only 38% of overall hippocampal length. In contrast, the PrS extended throughout the hippocampal head and body, and thus spans almost twice the length of PaS (32.6 mm/76.5%). Only a single case showing PrS tissue near the most posterior hippocampal boundary.

Based on this, a practical recommendation for in vivo segmentation would be to segment only the SUB in the most posterior slices of the hippocampus, given its consistent presence across cases (and datasets) in the tail region – with the important caveat that there is no unified definition of the tail. Regardless, our data on the disappearance of the PrS and PaS can help guide the posterior boundary of each region in segmentation protocols. For example, the PrS may reasonably be segmented across the anterior-to-mid portion of the hippocampus, corresponding to approximately three quarters of its total length.

### 3.4. Subicular complex boundary locations in coronal slices along the hippocampal body

*Aim:* A key question for potential in vivo MRI studies of subicular complex structure and/or function is determining the medial-lateral border placement for PrS/PaS and the proportion of these structures relative to the SUB when viewed on the coronal place (for a discussion of this see, Read et al., 2024; Dalton et al, 2017). Here, we aim to investigate the location of the PrS and PaS borders with respect to the length of the subicular complex and whether the position of these borders varies along the long axis of the hippocampal body.

*Approach:* Of the 17 available cases in the 2024 dataset, four were excluded due to missing segmentations or sections. The location of the PrS-SUB, and the PaS-PrS border, was measured relative to the width of the subicular complex on the coronal plane (Figure 5), that is, the SUB-CA1 border to the medial PaS border with the medial temporal lobe cortex. This was performed in five 10% increments (of total hippocampal length) from the uncal apex in each case, which corresponded to average increments of 2.6 mm (± 0.19 mm). As PaS is only present for a part of the hippocampal body (see Section 3.3), the slice selection was focused on this particular region to capture all subicular complex regions. To extract the length of each substructure in the coronal place, lines were drawn that bisected the cortex of each region. These were initiated on the CA1-SUB border and then extended to the PrS-SUB boundary, and then to the most medial PaS boundary. To capture the curvature of these regions in the coronal plane, lines were initiated from both regional borders (e.g., Figure 5a-e), as well as each major inflection point in the curvature (see Figure 5d).

**Figure 5.**
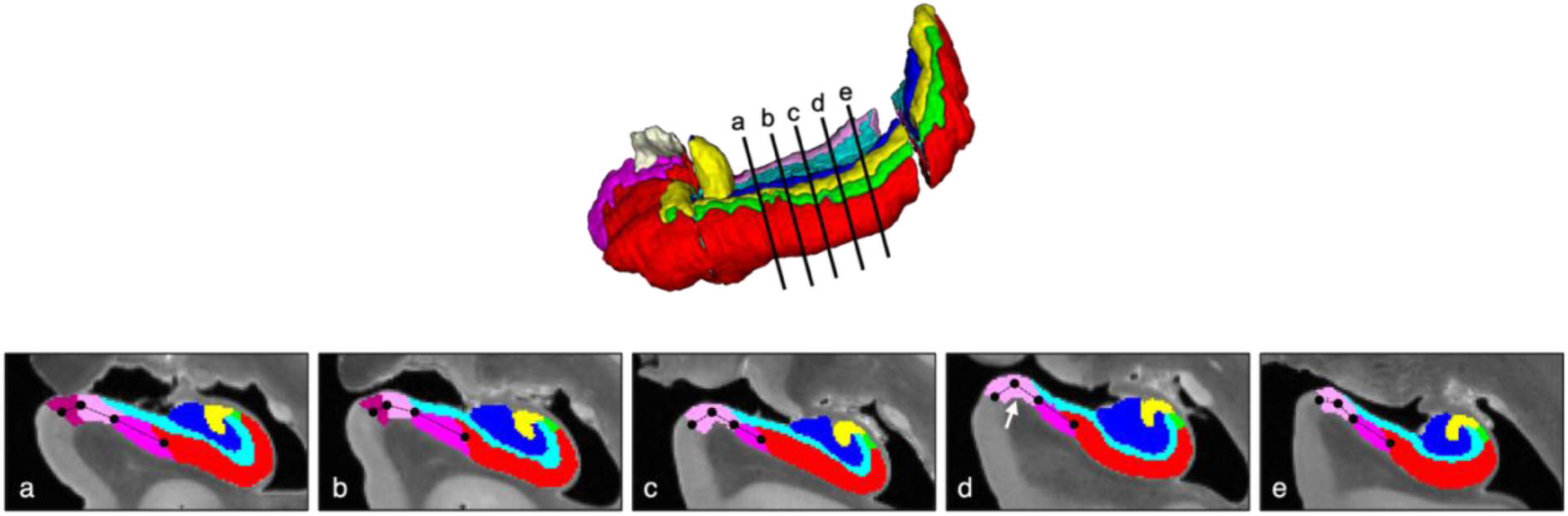
Approach for measuring the length and relative proportion of each subicular complex region in coronal slices along the hippocampal body. (a) Five slices were selected from the hippocampal body, which were sampled in 10% increments (of total hippocampal length) from the uncal apex. (b) To extract the length of each substructure in the coronal plane, lines were drawn that bisected the cortex of each region (in ITK-SNAP). These were initiated on the lateral border of the SUB and then extended to the PrS-SUB boundary, and then to the most medial PaS boundary. To capture the curvature of these regions in the coronal plane, lines were initiated from both regional borders (e.g., Figure 5a-e), as well as each major inflection point in the curvature (see Figure 5d).

*Results:* Consistent with the disappearance data reported in Section 3.3, the PaS was present throughout the hippocampal body in only three cases (CNDR14, CNDR20, CNDR23), with an average presence of 3.4 (±1.6) slices across cases. The SUB and PrS were observable in each slice increment. Overall, the SUB and PaS constitute the largest (*M* = 46.13%) and smallest (*M* = 15.33%) proportions of the subicular complex, respectively (see Table 4). In the cases where the PaS disappears, the relative proportion of the PrS increases accordingly (see Figure 6), which contributes to the augmented variation observed along the posterior axis.

**Figure 6.**
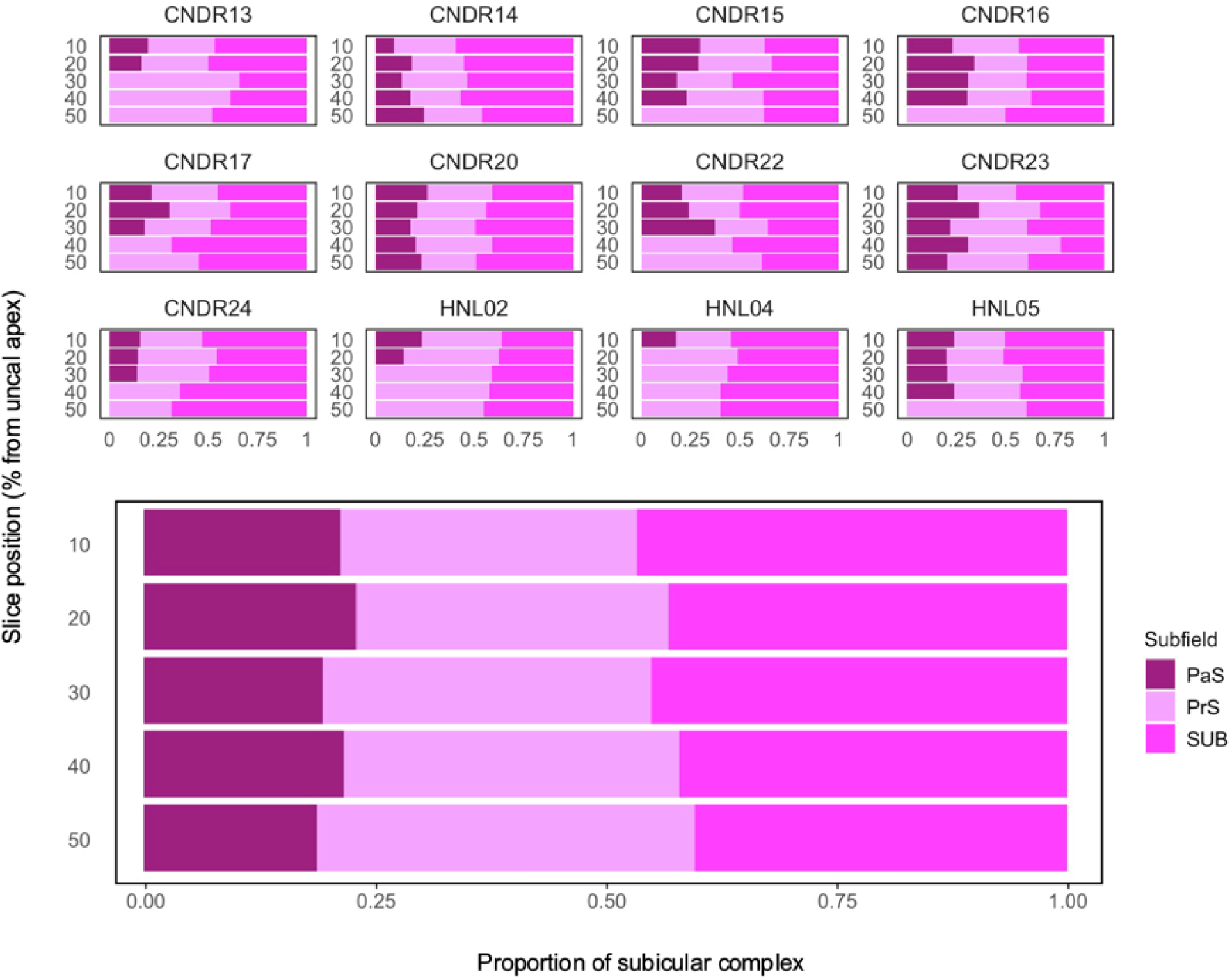
Mean length of subfields of the subicular complex as a percentage of the full subicular complex medial-lateral width for five coronal slices (a) as well as for each individual case (b). SUB = subiculum, PrS = presubiculum, PaS = parasubiculum.

**Table 4.** Percentage composition of the subicular complex in the coronal plane across different positions along the hippocampal body, showing the proportion comprised of the subiculum (SUB), presubiculum (PrS), and parasubiculum (PaS).

| **Position**<br>**along the**<br>**long axis**<br>**(%)** | **PaS** |  | **PrS** |  | **SUB** |  |
| --- | --- | --- | --- | --- | --- | --- |
|  | ***M*** | ***SD*** | ***M*** | ***SD*** | ***M*** | ***SD*** |
| **10** | 21.31 | 5.41 | 32.10 | 3.69 | 46.59 | 6.99 |
| **20** | 21.52 | 10.14 | 34.46 | 7.83 | 44.02 | 7.53 |
| **30** | 15.94 | 11.16 | 38.93 | 12.07 | 45.13 | 16.86 |
| **40** | 12.18 | 5.31 | 40.76 | 10.66 | 47.06 | 13.55 |
| **50** | 5.68 | 2.25 | 46.49 | 12.52 | 47.83 | 17.07 |
| ***Overall*** | 15.33 |  | 38.55 |  | 46.13 |  |

*M* = Mean in percent, *SD* = Standard deviation.

The medial-lateral border of the SUB and PaS appeared to be the most consistent throughout the hippocampal body, occupying between 46.6%-47.83% of the subicular complex. Thus, non-subicular proper regions were found to constitute more than half of the subicular complex length when viewed in the coronal plane. As the PaS disappears, the PrS absorbs that proportion of the subicular complex. Overall, the PrS and PaS together occupy between 37.5%-50.1% of the subicular complex, with the PrS occupying around 15.1% of that space. The PrS-PaS boundary showed no consistent evidence of becoming increasingly medial or lateral along the anterior-posterior axis. However, both inter- and intra-individual variations were observed. Some cases had a consistent boundary location throughout the hippocampal body (e.g. CNDR20 and CNDR24) while the border shifted laterally along the posterior axis or fluctuated in others (e.g. CNDR22 and CNDR23, respectively).

*Discussion*: These analyses yielded several key observations. The PaS consistently occupied the smallest proportion of the subicular complex, followed by PrS, and then finally SUB – a finding that is consistent with the anatomical literature in humans and monkeys (Ding, 2013). The most notable finding is that the medial-lateral boundary between PrS and SUB is located, with relative consistency across cases and slices, near the halfway point of the subicular complex, rarely exceeding (reaching *M* = 50.9%). Given this, a simple recommendation when segmenting PrS-PaS in the hippocampal body would be draw a subicular complex region that extends from the CA1 to the superior border of the entorhinal and parahippocampal cortices and then assign the most medial half of this region to PrS-PaS (with the caveat that PaS may not be presented in the posterior section of the hippocampal body). When considered alongside the disappearance data presented in Section 3.3, segmentation of a combined PrS-PaS region could cease around 8 mm anterior to the hippocampal tail endpoint (e.g., 4 x 2 mm slices), or at the emergence of the isthmus, at which point the PrS becomes contiguous with retrosplenial cortex (Barbas & Platt, 1995; Berger et al, 1997).

### 3.5. Location of CA3 relative to hippocampal head digitations

*Aim:* Hippocampal subfield segmentation protocols using in vivo MRI generally employ landmark-based heuristics for guiding border placement (Wisse et al, 2017; Canada et al, 2024; Olsen et al, 2019). The distinctiveness of the digitations in the hippocampal head may be one such useful landmark for segmentation, as they can be easily distinguished in vivo. Here we aimed to characterize the appearance of anterior CA3 relative to digitations in the hippocampal head in the 2024 dataset, similar to analyses performed in de Flores et al. (2019) in the 2019 dataset.

*Approach:* The anterior portion of CA3 was available in 11 cases in the 2024 dataset. The position of CA3 relative to the head digitations was taken as the most anterior coronal slice where CA3 appears in the superior strip of CA, that is, above the DG or SRLM. Unlike in de Flores et al. (2019), the most anterior CA2 was rarely present at the level of the head digitations but rather appeared more posterior, and thus not analyzed here. The most anterior CA1 appeared anterior to the start of the digitations and was not assessed for that reason. The determination of CA3 being classified as lateral, medial, or midpoint was given by where the bulk of CA3 exists (Figure 7). For example, in Figure 7a all of CA3 is medial to the midpoint. In Figure 7b CA3 exists around the midpoint in equal lateral and medial parts and is thus classified as midpoint.

**Figure 7.**
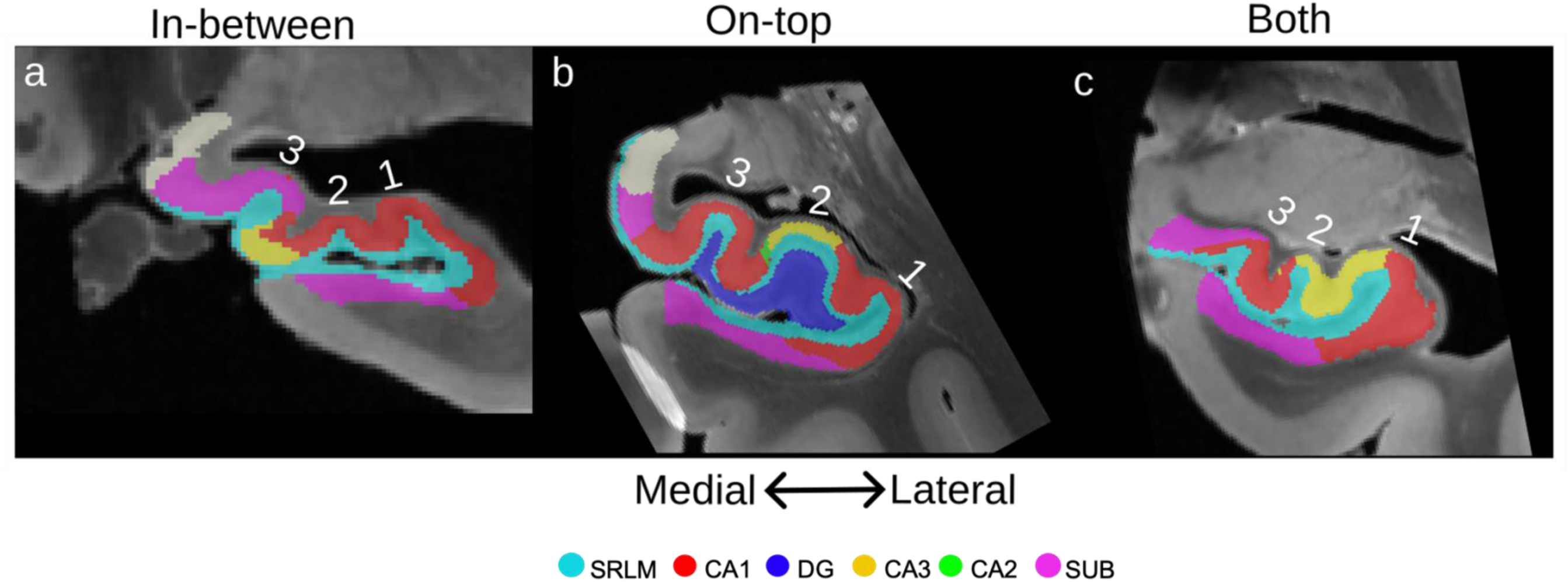
Depiction of the most anterior CA3 (yellow) relative to the digitations of the hippocampal head. Digitations are numbered from lateral to medial. (a) CA3 appears in-between the digitations and presents more medially relative to the whole width of the hippocampus. (b) CA3 appears on-top of a digitation and presents at the midpoint between medial and lateral. (c) CA3 appears both on-top and in-between the digitations and presents more laterally relative to the whole width of the hippocampus. SRLM = strata radiatum lacunosum moleculare, CA = cornu ammonis, DG = dentate gyrus, SUB = subiculum.

*Results:* The majority of cases had 2 (36%) or 3 (45%) digitations (Table 5). Anterior CA3 largely appeared in-between the digitations (64%; Figure 7a) followed by on-top of a digitation (18%; Figure 7b) and both on-top and in-between (18%; Figure 7c). CA3 presented equally in medial (36%; Figure 7a) and lateral (36%; Figure 7c) positions, followed by at the midpoint between medial and lateral (28%; Figure 7b).

**Table 5.** Quantification of the number of digitations and the location of anterior CA3 relative to the digitations.

|  |  |
| --- | --- |
| **Number of digitations (mean)**<br>- 1 digitation (n [% total])<br>- 2 digitations (n [% total])<br>- 3 digitations (n [% total])<br>- 4 digitations (n [% total]) | **2.5**<br>- 1 [9%]<br>- 4 [36%]<br>- 5 [45%]<br>- 1 [9%] |
| **Location of anterior CA3**<br>- On-top of digitation (n [% total])<br>- In-between digitations (n [% total])<br>- Both on-top and in-between (n [% total]) | - 2 [18%]<br>- 7 [64%]<br>- 2 [18%] |
| **Position of anterior CA3 relative to digitations**<br>- Lateral (n [% total])<br>- Medial (n [% total])<br>- Midpoint (n [% total]) | - 4 [36%]<br>- 4 [36%]<br>- 3 [28%] |

*Discussion:* The mean number (2.54) and range (1-4) of head digitations found here aligns with that the mean of 2.88 found in de Flores et al. (2019) and of mean of 2.46 found in Gertz et al. (1972). While there is considerable inter-case variability, there does seem to be a pattern of CA3 appearing in-between the digitations, which aligns with previous research (de Flores et al., 2019). However, we found that the appearance of anterior CA3 varied almost equally between lateral, medial, and midpoint positions. As well, CA2 appeared much more posterior to the head digitations here, in contrast to the segmentations in de Flores et al. (2019), suggesting that there can be great variability in the anterior starting point of CA2 between neuroanatomy laboratories or datasets. Thus, the head digitations are unlikely to be a useful landmark for subfield segmentation since CA3 cannot be reliably located apart from a general rule of appearing in-between the digitations.

### 3.6. Subfield borders in the body of the hippocampus relative to a volume proportion of the SRLM around the DG

*Aim:* The development of methods to segment the hippocampal subfields using MRI have generally relied on a combination of anatomical landmarks and geometric rules (de Flores et al., 2019; Steve et al., 2017; Yushkevich et al., 2015). The SRLM and the DG are two such landmarks. The location of the subfield borders has been shown to be at consistent relative curvilinear lengths along the SRLM (de Flores et al., 2019; Steve et al., 2017). The aim of this section is to investigate if the subfields (specifically, CA1, CA2, and CA3) project onto a consistent relative volume proportion of the SRLM around the DG throughout the hippocampal body in the 2024 dataset, and if such a measure may be useful for guiding in vivo segmentations.

*Approach:* The SRLM label was replaced in 15 cases based on a projection of the CA1, CA2, and CA3 borders onto its volume (Figure 8). This was done in 10% increments (up to 50% - still within the hippocampal body) posterior from the uncal apex, similar as in section 3.4 (Figure 5). Each 10% increment corresponded to a 2.6 mm distance on average. Only the portion of the SRLM that is lateral to the most medial point of the DG was considered (Figure 8), given that this section can usually be reliably identified on in vivo MRI. The proportion of total SRLM voxels that were replaced by each subfield was then calculated as the number of SRLM voxels (lateral to the medial DG) replaced by a given subfield projection divided by the total number of SRLM voxels (lateral to the medial DG) for that slice (Figure 8). The bottom row of Figure 8 depicts a case where the subiculum (pink)/CA1 (red) border occurs lateral of the medial DG border. Although the whole SRLM lateral to the DG is used for the calculation, in such a case the projection of CA1 onto the SRLM was taken as the middle of the SUB-CA1 border. Note that in these cases the CA1, CA2, and CA3 SRLM proportion do not sum to unity.

**Figure 8.**
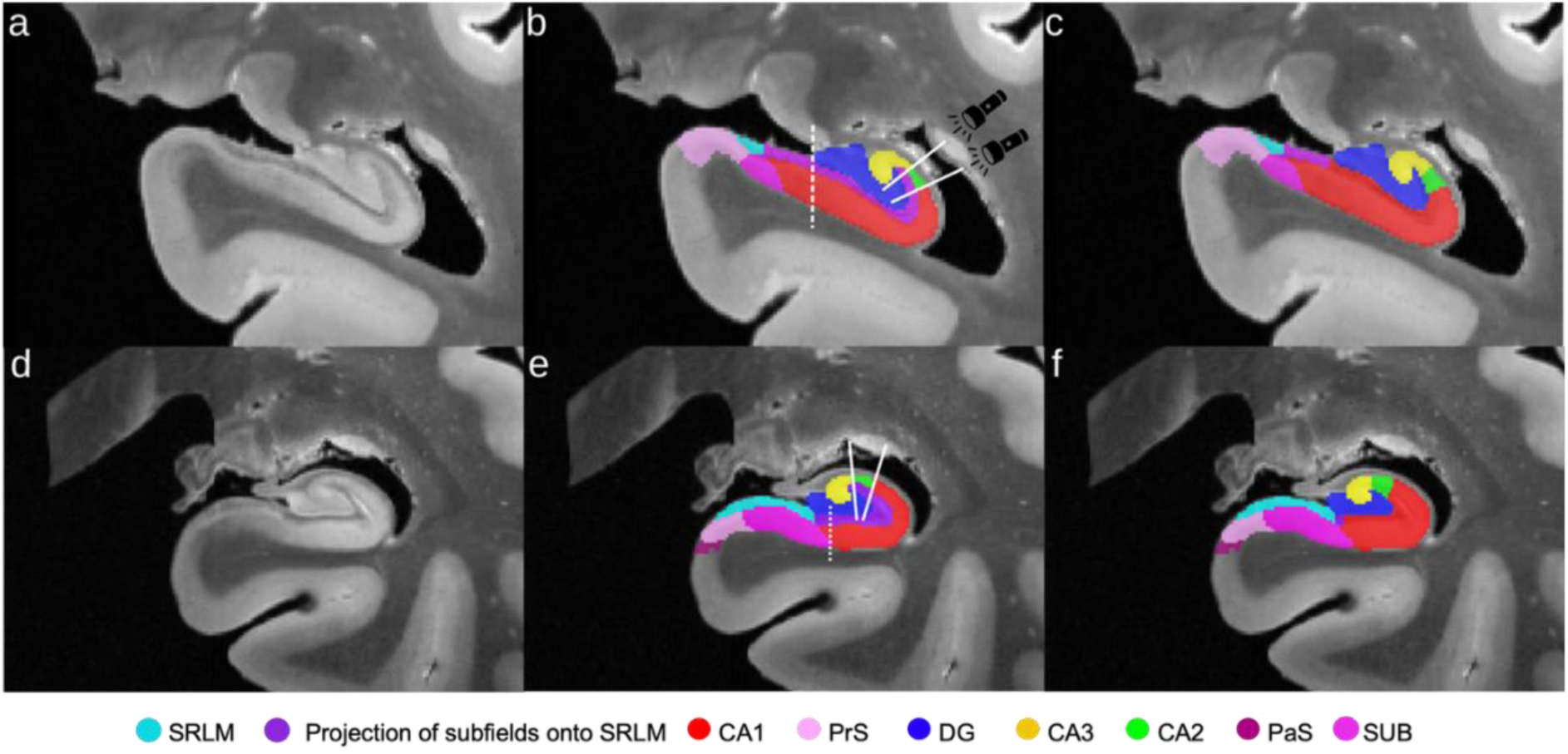
Flashlight analogy for projection of subfields onto SRLM (purple), where the “beam” of light is projected across the subfield borders onto the SRLM (b) which is re-labelled proportionally (c). Only the portion of the SRLM that is lateral to the most medial point of the DG (dark blue) is considered (white dashed line in b). The bottom row depicts a case where the subiculum (pink)/CA1 (red) border occurs lateral of the medial DG border. Although the whole SRLM medial to the DG is used for the calculation, in such a case the middle of the subiculum/CA1 border (dotted line in e) is taken for the projection seen in (f) to determine the proportion of the SRLM encompassed by CA1. Note the small unlabeled portion of the SRLM in (f), thus cases exist when CA1, CA2, and CA3 do not encompass the whole volume of the SRLM that is lateral to the medial DG. SRLM = strata radiatum lacunosum moleculare, CA = cornu ammonis, PrS = presubiculum, DG = dentate gyrus, PaS = parasubiculum, SUB = subiculum.

*Results:* On average, CA1 encompasses around 83.5% of the SRLM lateral to the medial DG, CA2 encompasses around 9.2%, and CA3 encompasses around 4.0% (Figure 9; Supplementary Table 3). As well, the percent of the SRLM encompassed by the three CA subfields did not appear to change much for any subfield from anterior to posterior portions of the hippocampal body. The percent of CA1 around the SRLM is more variable than CA2 or CA3, as can be seen from the larger interquartile range. Given that this is a measure of volume proportion around the SRLM, we also looked to compare these values to subfield volume at the same slices as above (Supplementary Figure 4). The measures of subfield volume tend to be more variable than the SRLM volume proportion measure described here, particularly for CA1.

**Figure 9.**
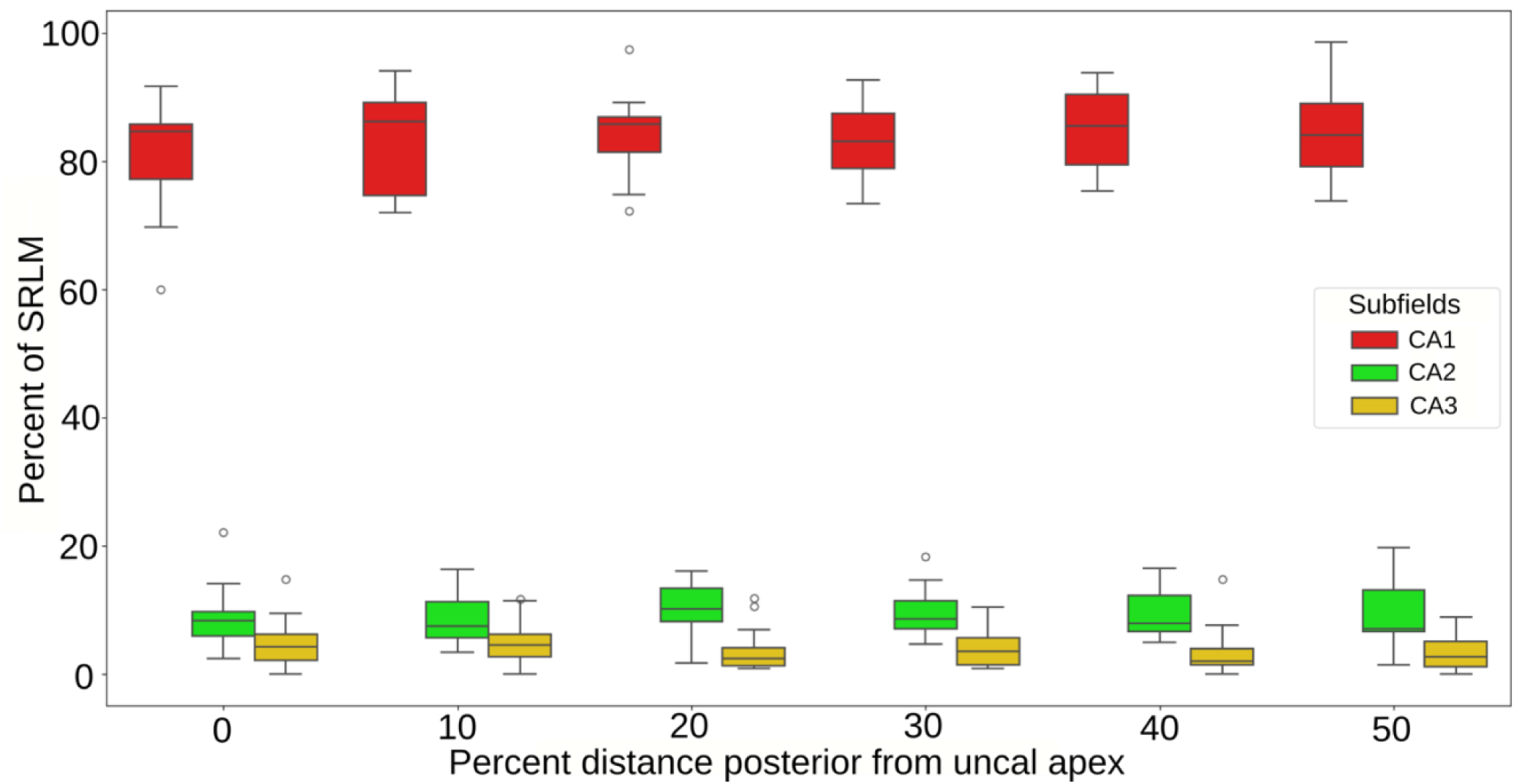
Proportion of the SRLM located lateral to the most medial point of the DG that CA1, CA2, and CA3 encompasses along the longitudinal axis of the hippocampal body. CA = cornu ammonis.

*Discussion:* The SRLM proportionality measure derived here was examined to see if it could provide a useful benchmark for evaluating in vivo segmentations. However, the measure does not explicitly quantify subfield border position, rather it quantifies the relative size of the subfields. It appears to be much more consistent than the typical measure of subfield volume, which has much more inter-individual variability (Supplementary Figure 4). In Section 3.5 of de Flores et al. (2019) and in Steve et al. (2017) the distance (in mm) of each subfield border relative to the SRLM was calculated by tracing a line which follows the curvature of the SRLM from the medial point of the DG (same starting point as in the current study). On average, de Flores et al. (2019) found that the SUB-CA1 border occurred at 33% along the curved length of the SRLM from the most medial point of the DG in a lateral direction, and the CA1-CA2 border occurred at 80% along the SRLM length. Thus, based on these measures, CA1 can be said to occupy 47% (80% - 33%) of the curved SRLM length. Using the same reasoning, Steve et al. (2017) found that CA1 occupied 68.7% of the curved SRLM length on average. In the current study we found that CA1 encompassed on average 83.5% of the SRLM volume lateral to the most medial point of the DG. Following the same reasoning, Supplementary Table 4 reports the percentages for CA2 and CA3 between de Flores et al. (2019), Steve et al. (2017), and the current study. The percentage of CA2 around the SRLM of the current study closely matches that found in de Flores et al. (2019; 9.4 and 10.6% respectively), while for CA3 it is closer to the estimate found in Steve et al. (2017; 4 and 2.5% respectively). The difference in the proportion of the subfields encompassing the SLRM between the current study and de Flores et al. (2019) and Steve et al. (2017) may be due to a few factors. Firstly, the protocols for defining the cytoarchitectonic boundaries are not the same, such that the location of the subfield borders may differ systematically. Secondly, the thickness of the SRLM, which is relevant for the current proportion measure, but is not relevant in de Flores et al. (2019) and Steve et al. (2017), may not be consistent along its length in some subjects. Thus, the subfield borders could be in the same place, but if the SRLM were thicker around CA1, for example, this would result in different proportions from de Flores et al. (2019) and Steve et al. (2017). While the method in de Flores et al. (2019) and Steve et al. (2017) can provide the location of each subfield border exactly in millimeters, it requires the use of external software not used for subfield segmentation and is quite labor intensive, making it difficult to apply. The current method may be useful as an in vivo segmentation rule, particularly for CA2 and CA3. While our calculated proportion of CA1 was further from that in de Flores et al. (2019) and Steve et al. (2017), our calculated proportion of CA2 and CA3 was relatively consistent.

### 3.7. Hippocampal length as a function of demographic factors and factors specific to postmortem studies

*Aim:* Segmentation protocols for the whole hippocampus, hippocampal subfields, and surrounding medial temporal lobe cortices often define anatomical boundaries using a fixed number of slices from a certain landmark, or a percentage of total length of the hippocampus or hippocampal head (Moore et al., 2014 Berron et al., 2017; see also rule proposals in Sections 3.1-3.3 above). The question is, though, whether hippocampal length is subject to change in relation to demographic factors or diagnostic status, which could then bias the segmentation protocol. For example, if a certain segmentation rule has to be applied for a specific number of slices, this would lead to a different result in short versus long hippocampi and could particularly bias the results if hippocampal length is related to a clinical variable of interest. We therefore aimed to investigate the associations of age, dementia status, and sex with hippocampal head length and total hippocampal length. Additionally, because we utilized postmortem MRI, we aimed to assess the potential confounding factors of fixation time and postmortem interval (time between death and autopsy) in this type of study by examining their associations with hippocampal head length and total hippocampal length. Finally, we investigated the ratio between hippocampal head length and total length to see whether the head or posterior part of the hippocampus could be disproportionately affected by any of the variables of interest.

*Approach:* The hippocampal length measures, that is, head length, total length, and head-to-total length ratio, were obtained from cases across both atlases. Head length, defined as the distance from the most anterior tip of the hippocampus to the most posterior tip of the uncal apex, was measured in all 26 cases. Total length, defined as the distance from the most anterior tip of the hippocampus to the most posterior slice of the hippocampus, was measured in 24 out of 26 cases. In two cases (HNL01 and HNL03), the most posterior slice could not be determined, which prevented the calculation of total length. Consequently, the ratio was calculated for 24 cases by dividing the head length by the total length and expressing the result as a percentage. The determination of the most anterior and posterior slices of the hippocampus has been described in sections 3.1 and 3.2.

Mann–Whitney U tests were conducted to assess the associations between the hippocampal length measures and dementia status and sex. Spearman’s rank correlations were used to examine the associations between the hippocampal length measures and age, fixation time, and postmortem interval. Analyses regarding dementia status included both cases with and without dementia, while tests for sex, age, fixation time, and postmortem interval were restricted to cases without dementia. The Benjamini–Hochberg procedure was applied separately to each family of length analyses (head, total, and ratio) to control the false discovery rate (FDR). A significance level of 0.05 was set for all analyses, which were performed using IBM SPSS Statistics (version 29).

*Results:* Dementia status was not significantly associated with hippocampal head length (U = 46.50, p = 0.54), total hippocampal length (U = 52.50, p = 0.73) or the ratio (U = 35.00, p = 0.33; Table 6), although hippocampal head and total length were about 1 mm shorter in cases with dementia compared to those without dementia (Supplementary Table 5). Similarly, sex was not significantly associated with hippocampal head length (U = 5.50, p = 0.657), total hippocampal length (U = 3.00, p = 0.73), or the ratio (U = 4.00, p = 0.67). Qualitatively, women had ∼1 mm shorter hippocampal head length and ∼3 mm shorter total hippocampal length. Age was also not significantly correlated with hippocampal head length (r_s_ = -0.33, p = 0.66), total hippocampal length (r_s_ = -0.38, p = 0.73) or the ratio (r_s_ = -0.78, p = 0.21), although the correlation coefficients are relatively high.

**Table 6.** Association of hippocampal length measures with dementia status, sex, age, fixation time, and postmortem interval. All *p*-values, except for that of the correlation between head length and total length, were FDR-corrected.

|  | Dementia status |  | Sex |  | Age |  | Fixation time |  | Postmortem interval |  | Total length |  |
| --- | --- | --- | --- | --- | --- | --- | --- | --- | --- | --- | --- | --- |
| | $U$ | $p$ | $U$ | $p$ | $r_s$ | $p$ | $r_s$ | $p$ | $r_s$ | $p$ | $r_s$ | $p$ |
| Head length | 46.50 | 0.535 | 5.50 | 0.657 | -0.33 | 0.657 | 0.06 | 0.881 | 0.22 | 0.722 | 0.73 | <0.001 |
| Total length | 52.50 | 0.728 | 3.00 | 0.728 | -0.38 | 0.728 | -0.23 | 0.728 | -0.16 | 0.728 |  |  |
| Ratio | 35.00 | 0.325 | 4.00 | 0.67 | -0.78 | 0.205 | -0.20 | 0.67 | 0.52 | 0.382 |  |  |

FDR = false discovery rate, $U$ = Mann–Whitney $U$ test statistic, $p$ = $p$ -value, $r_s$ = Spearman's rank correlation coefficient.

Fixation time and postmortem interval were also not associated with hippocampal head length (r_s_ = 0.06, p = 0.88; r_s_ = 0.22, p = 0.72), total hippocampal length (r_s_ = -0.23, p = 0.73; r_s_ = -0.16, p = 0.73) or the ratio (r_s_ = -0.20, p = 0.67; r_s_ = 0.52, p = 0.38).

Finally, a strong positive correlation was found between head length and total length (r_s_ = 0.73, p < 0.001; Supplementary Figure 5).

*Discussion:* We found no significant association of hippocampal length (as measured by head length, total length, and head-to-total length ratio) with dementia status, sex, or age. Variables specific to postmortem imaging, such as fixation time and PMI, suggest that the findings in the current section are directly applicable to in vivo MRI. However, there are several key caveats. First, the small sample size, particularly the limited number of controls, restricted our ability to detect more subtle differences between groups and to examine inter-case differences. Indeed, there were some moderate-to-strong effect sizes observed (e.g., between all length measures and age, and between ratio and postmortem interval) that would require much larger samples to extract. Second, there was substantial heterogeneity within the dementia group, which included cases with various subtypes of dementia, potentially affecting the hippocampus differently. This variation could have reduced the sensitivity required to detect the effects of dementia on hippocampal length. Finally, five out of nine control specimens in the head length analysis and three out of seven in the total length and ratio analyses had unknown clinical statuses. Although it is assumed that these controls did not suffer from dementia, it is possible that some latent pathology in the control group masked any potential differences. Larger-scale replication studies are necessary to confirm our results. The demographic factors of age, sex, and dementia status especially could easily be replicated in a larger in vivo study. Additionally, it could be of interest to explore the potential association of other diagnoses with hippocampal length measures.

### 3.8. Relative location of SUB-CA1 border in coronal slices in the hippocampal body as a function of demographic factors and factors specific to postmortem studies

*Aim:* Several challenges arise when identifying the SUB-CA1 border in MRI images. There is considerable interindividual variability in the border location (Zeineh et al, 2015) as well as differences in the cytoarchitectonic definition of this border between different neuroanatomy laboratories (Wuestefeld et al, 2024). Not surprisingly, the variability in the SUB-CA1 border is reflected in the segmentation protocols for in vivo MRI (Yushkevich et al, 2015). Some interindividual variation may be related to demographic or clinical variables, such as dementia. de Flores et al (2019) found that dementia cases, but not non-dementia cases, exhibited a medial shift in the location of the SUB-CA1 border progressing posteriorly along the longitudinal axis. Our aim in this section was to see if this finding persists when extending the dataset, as well as to investigate the relationship between the SUB-CA1 border location and demographic variables age and sex and factors specific to postmortem imaging, i.e. fixation time and postmortem interval, to see if interindividual variability is driven by these parameters.

*Approach:* One case was excluded from each dataset, with a total of 24 cases used for the analyses. The method for measuring the relative position of the SUB-CA1 border follows that of de Flores et al (2019), see Figure 10. The location of the SUB-CA1 border was measured in reference to the combined width of the DG and cornu ammonis structures or lateral portion of the hippocampal body. This was operationalized as the distance between the most medial anchor point and the edge of the most lateral CA1 voxel, and the DG. The medial anchor point itself was defined by first identifying the most medial edge of CA3 or DG (whichever subfield had the most medial voxel), and then by drawing a straight, vertical line to the SUB or CA1 (Figure 10a; Supplementary Figure 6). The point where this line touched CA1 or SUB was pinpointed as the most medial anchor point (de Flores et al, 2019). A vertical line was drawn from the SUB-CA1 border (from its midpoint, if at an angle) to the superior edge of the DG, and the location of the SUB-CA1 border was then quantified as the distance from the medial anchor point to this line. See Supplementary methods for section 3.8 for more detail.

**Figure 10.**
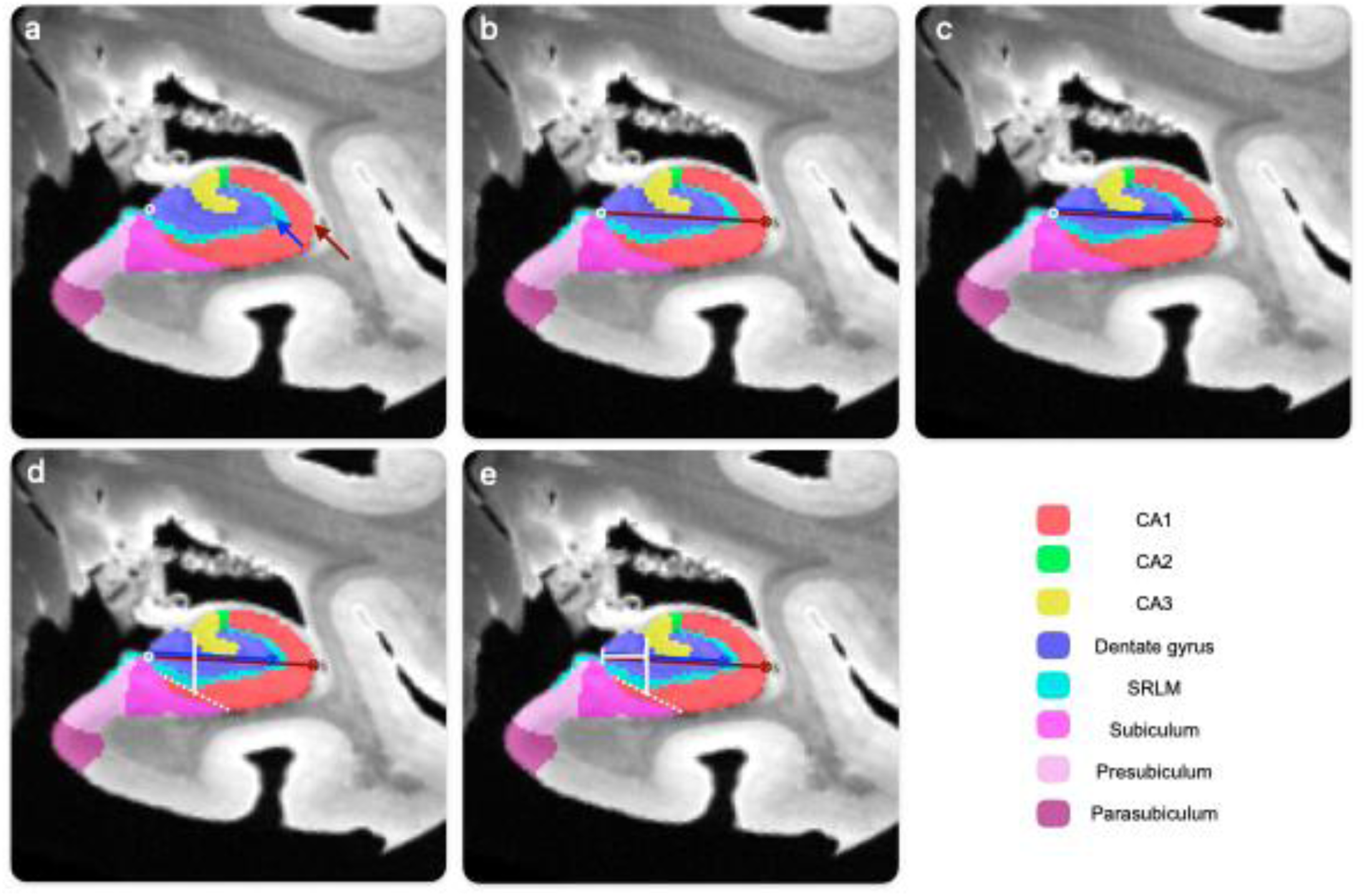
Approach to measure the SUB-CA1 border location relative to the lateral portion of the hippocampus in the hippocampal body in an example case. a) First, the medial anchor point, i.e. the most medial voxel at which the DG meets the SRLM/subicular complex, was identified (white circle) as well as the most lateral edge of the CA1 and DG (blue and red arrows). b) Second, the full hippocampal width was measured by taking the distance between the medial anchor point and the lateral edge of the CA1 (red line). c) Third, the width of the DG was measured by taking the distance between the medial anchor point and the lateral edge of the DG (blue line). d) Fourth, the SUB-CA1 border (dashed white line) was identified and a vertical line from the midpoint of this border (solid white line) was drawn. e) Fifth, the distance from the medial anchor point to the vertical midpoint line was measures (orange line). The relative position of the SUB-CA1 is expressed as percentage of the reference width, i.e. either the full hippocampal width or the DG width. SUB = subiculum, DG = dentate gyrus, CA = cornu ammonis, SRLM = strata radiatum lacunosum moleculare.

These measurements were taken at two points, one slice posterior to most posterior tip of the uncal apex (here after referred to as 0mm) and once 20 mm posterior to the uncal apex itself. Slices within a +/-3 mm range were included to circumvent incomplete segmentation coverage. If no appropriate slice could be found within this range, the case was excluded for this position. The measurements were redone for the 2019 dataset to limit inter-rater variability effects and allow for grouping of cases across the two datasets.

The statistical analyses were performed separately for the hippocampal and DG width and both positions in the hippocampal body. The statistical analyses were the same as in section 3.7, but were performed using R (R Core Team, 2025).

*Results:* Overall, there appears to be a medial shift of the SUB-CA1 border along the anterior-posterior axis (see Table 7). When combining the 2019 and 2024 dataset, the mean relative position of the of the SUB-CA1 border at 0 mm from the uncus is 25% (SD = 25.3) of the full hippocampal width and 32.9% (SD = 32.6) of the DG width. At 20 mm, the relative position has shifted to 0.9% (SD = 33.3) of the full hippocampal width and 1% (SD = 46.7) of the DG width. This shift was statistically different for both the full hippocampal width (*U* = 272, p<0.001) and the DG width (*U* = 270, p<0.001). Furthermore, there is more variability in the location of the SUB-CA1 border in the posterior compared to the anterior hippocampus as indicated by a higher standard deviation. These results are comparable to de Flores et al (2019), but the medial shift is slightly larger than previously observed (Supplementary Table 6). Comparing the two datasets, the 2024 dataset tended to have more medial SUB-CA1 border than the 2019 dataset, but it is important to note that there is also substantially more variability in the 2024 dataset (Table 7).

**Table 7.** Relative position of the SUB-CA1 border of the hippocampal and DG width, measured from medial (0%) to lateral (100%).

|  | **2019<sup>a</sup>** |  | **2024** |  | **2019 + 2024** |  |
| --- | --- | --- | --- | --- | --- | --- |
| **Reference** | **0 mm**<br>**mean**<br>**(SD)** | **20 mm**<br>**mean (SD)** | **0 mm mean**<br>**(SD)** | **20 mm mean**<br>**(SD)** | **0 mm mean**<br>**(SD)** | **20 mm mean**<br>**(SD)** |
| **% Of CA+DG**<br>**width** | 43.6<br>(15.1) | 17.6 (18.2) | 14.2 (25) | -7.39 (36.4) | 25 (25.3) | 0.9 (33.3) |
| **% of DG**<br>**width** | 56.7<br>(17.9) | 22.6 (23.4) | 18.6 (32.5) | -9.86 (52.1) | 32.9 (32.6) | 1 (46.7) |

<sup>a</sup>New rater; for comparison with rater from De Flores et al (2019), see Supplementary Table 5. All cases excluding NDRI02 and CNDR19 (missing segmentations). Provided means are in percent. SUB = subiculum, CA = cornu ammonis, DG = dentate gyrus.

The relationships between location of the SUB-CA1 border, whether relative to hippocampal or DG width, at 0mm or 20mm from the uncus and demographic or factors related to post-mortem imaging were not significant (Table 8). Overall, non-dementia cases qualitatively tended to have a more medial SUB-CA1 border location throughout the hippocampus (see Supplementary Table 7). This was driven by the 2024 dataset, in which the border progressed more medial than the most medial designated anchorpoint at 20 mm, as the reverse was observed in the 2019 dataset. Moreover, relatively strong though not significant associations were observed of age and fixation time and the SUB-CA1 border location especially at the 20 mm location.

**Table 8.** The relationship between the relative SUB-CA1 border location and demographic/imaging-specific factors from both datasets.

| Distance from the uncus | Reference width | Dementia<sup>a</sup> |  | Sex<sup>b</sup> |  | Age<sup>b</sup> |  | Fixation days<sup>b</sup> |  | PMI<sup>b</sup> |  |
| --- | --- | --- | --- | --- | --- | --- | --- | --- | --- | --- | --- |
|  |  | *U* | *p* | *U* | *p* | *r<sub>s</sub>* | *p* | *r<sub>s</sub>* | *P* | *r<sub>s</sub>* | *p* |
| 0mm | CA+DG | 85 | 0.35 | 10 | 0.69 | 0.52 | 0.35 | 0.62 | 0.35 | -0.36 | 0.48 |
|  | DG | 84 | 0.35 | 10 | 0.69 | 0.49 | 0.35 | 0.62 | 0.35 | -0.36 | 0.48 |
| 20mm | CA+DG | 69 | 0.23 | 8 | >0.99 | 0.74 | 0.12 | 0.74 | 0.12 | -0.22 | 0.76 |
|  | DG | 71 | 0.23 | 8 | >0.99 | 0.74 | 0.12 | 0.74 | 0.12 | -0.22 | 0.76 |

*U* = Mann–Whitney U test statistic, *p* = *p*-value, *r<sub>s</sub>* = Spearman’s rank correlation coefficient. All *p*-values are FDR-corrected. <sup>a</sup>N=24. <sup>b</sup>N=8; only non-dementia cases were included. SUB = subiculum, CA = cornu ammonis, DG = dentate gyrus.

Furthermore, two influential outliers were identified by comparing box-plots and using the R package rstatix (Kassambara, 2023). Analyses were rerun without these outliers, but yielded only minor differences (<1mm difference) and did not alter major.

*Discussion:* The inclusion of the 2024 dataset provided important insights into SUB-CA1 border delineation. A key finding was the medial shift in the position of the SUB-CA1 border along the anterior-posterior axis, consistent with the findings of de Flores et al (2019). This shift was present in both dementia and non-dementia cases, in both male and female cases. There also appeared to be greater variation in the location of the SUB-CA1 border in the posterior part of the hippocampus, which was also observed by de Flores et al (2019). There are some notable differences between the datasets, specifically markedly greater variation in the border location in the 2024 dataset, especially at 20mm, suggesting there is greater between-subject variability than previously reported.

A key difference between these findings and the findings of de Flores et al (2019) is that the medial shift along the anterior-posterior axis did not appear to be dementia-driven upon the inclusion of the 2024 dataset. Qualitative differences between dementia cases and non-dementia cases did not reach significance, in the combined dataset or the individual datasets, contrary to the findings of de Flores et al (2019). While inter-rater differences were non-significant, the current measurements for the 2019 datasets were more variable, potentially due to slight differences in measurement approaches. However, there were several 2024 cases where the SUB-CA1 border was notably more medial than the most medial anchorpoint, which did not feature in any of the 2019 cases. This subset of cases is not explainable by inter-rater discrepancies, but rather indicative of a heterogeneity in SUB-CA1 border location which is not attributable to dementia status, other demographic factors, or factors specific to postmortem studies alone, this warranting some caution in using a single segmentation rule for all cases.

While the sample size is relatively large for ex-vivo studies of the hippocampus, it is still limited for the purpose of drawing firm conclusions. For example, it is likely that the analyses presented are under-powered, especially with the great variability observed; one example of this is the relationship between age and the relative SUB-CA1 border location, which showed a strong but non-significant correlation. A small sample size is also more vulnerable to outliers, such as the cases in the 2024 dataset where the SUB-CA1 border was more medial than the anchor point, which reiterates the need for caution when attempting to disseminate segmentation rules from a limited number of cases. In sum, while some demographic and postmortem imaging factors were not related to the relative position of the SUB-CA1 border, effect sizes for age and fixation time were relatively large and warrant further research in larger postmortem samples. These findings warrant caution towards using the relative location of the SUB-CA1 border for in vivo segmentation as the findings may be affected by fixation time and age.

## 4. Discussion

This study builds upon the previous work of de Flores et al (2019) through the use of a larger postmortem MRI dataset that includes histology-based hippocampal subfield segmentations and detailed neuropathological information. This enabled further characterization of hippocampal subfield borders and organization, with direct relevance to segmentation approaches on in vivo MRI. The main findings of this study are as follows: a) hippocampal subfields appeared in a consistent order across most cases with some variability between CA2, CA3 and DG; b) the SUB and CA1 are present until the posterior tip of the hippocampal tail but there is greater variability in the CA3, DG and CA2; c) subicular complex subfields follow a consistent order of appearance with an inverted order for their disappearance; d) the PrS and PaS combined consistently occupy around half of the subicular complex on coronal slices in the hippocampal body, with some inter- and intraindividual variability on exact boundary locations; e) the location in which the CA3 appears varies considerably and hippocampal digitations are not a reliable landmark; f) SRLM proportionality offers a consistent approach for estimating CA2 and CA3 subfield borders; g) hippocampal length measures were not significantly related to demographic factors or factors specific to post-mortem imaging; and h) there is considerable variation in the SUB-CA1 border location but the source of this variation was not significantly related to diagnosis, demographic factors or factors specific to post-mortem studies. Certain findings from de Flores et al (2019) were replicated – for example, the order of appearance of CA1 and SUB, the presence of CA1 and SUB until the posterior tip of the hippocampus, the medial shift of the SUB-CA1 boundary in the hippocampal body, and the lack of association between the location of CA3 and the digitations. Other findings diverged from this earlier study, such as the CA3 appearing anterior to CA2 in the 2024 dataset, the presence of CA2, CA3 and DG in the posterior tip in the hippocampus, the association of the relative SUB-CA1 border with dementia status, and the relative proportion of CA2 and CA3 to SRLM volume.

A key novel contribution of the present study is the detailed description of how subicular complex regions are organised along the anterior-posterior axis of the hippocampus. The subicular complex, which includes SUB, PrS, and PaS, has been implicated in a range of brain functions, including episodic memory, scene processing, navigation, stress, anxiety, and reward processing (Aggleton & Christiansen, 2015; O’Mara et al, 2001; O’Mara et al, 2005). Its diverse functional roles, as well as its potential involvement in memory disorders, underscore the need for a more detailed understanding of its anatomical properties, particularly where histological evidence can aid in distinguishing its constituent regions in the human brain. Recent human MRI studies have highlighted uncertainty in determining whether functional activations or streamline terminations from diffusion tractography in the medial subicular complex correspond to SUB or to PrS/PaS (Read et al, 2024; Dalton et al, 2022; Hodgetts et al, 2017). By documenting the appearance and disappearance of subicular complex regions along the longitudinal axis of the hippocampal formation, both relative to each other and with respect to neighbouring subfields (e.g., CA1-3 and DG), our findings offer useful heuristics for interpreting statistical parametric neuroimaging maps. While this study provides practical guidelines for subdividing the subiculum in the hippocampal body (e.g., defining a boundary halfway between the subicular complex, entorhinal cortex and SUB-CA1 border), further work is needed to clarify the spatial arrangement of PrS and PaS in the hippocampal head. An important next step will be to integrate the observations presented here, collected across a range of cases, with existing segmentation schemes (Dalton et al, 2017; Hickling et al, 2024) to develop more reliable and valid anatomical delineation rules.

This study featured two post-mortem datasets totaling 26 cases, with and without dementia and within a wide age-range, which allowed for a comprehensive examination of hippocampal subfields characterization across a variety of pathological and demographic profiles. A key takeaway from our results is the importance of interindividual variation. While certain features, such as the order of subfield appearance and disappearance, exhibit consistent patterns, other aspects, like the proportions of subicular complex subfields, may demonstrate considerable variability between and within individuals. Including a greater diversity of cases has revealed that the source of this variation may be quite complex and not attributable to a single factor. It is likely that biological differences only represent part of this variability, with differences in neuroanatomical schools of thought and training augmenting inconsistencies (Wuestefeld et al, 2024). Thus, this study further underscores the need for consensus on anatomical landmarks and segmentation protocols.

Although this study included cases with dementia, the grouping of these cases into a single category might have masked diagnosis-specific variations in hippocampal subfield characteristics. While it not possible to characterize hippocampal subfield anatomy in relationship to specific neurodegenerative diseases, this pragmatic grouping may obscure nuances associated with different neuropathological and clinical diagnoses. For example, these results indicated that there was no significant difference in hippocampal length between dementia cases and non-dementia cases, which differs from the findings of other papers (e.g. Dawe et al., 2011). A more granular analysis may yield more specific insights, as even within-diagnosis, e.g. Alzheimer’s disease, there are systematic anatomical differences associated with disease progression. Lastly, due to the sample size, some inferential statistics may be underpowered, which limits the ability to draw definitive conclusions. The present paper is primarily exploratory in its approach, aiming to identify and contextualize broad patterns within the two datasets to inspire further lines of inquiry and stimulate the development of segmentation guidelines. Specifically, the relative strong effect sizes observed for age in relationship hippocampal length measures, but also the notable qualitative differences in hippocampal length measures between the sexes could easily be further investigated in large in vivo MRI studies. Unfortunately, some of the potential underpowered associations of demographics or postmortem imaging variables in relationship to the relative SUB-CA1 border are more difficult to investigate in larger or in vivo MRI samples, as histological validation is required.

This study does, however, provide both informative and actionable insights. One clear finding is that the order of appearance of certain subfields in the anterior hippocampus is remarkably consistent, as is the presence of specific subfields at the posterior tip. This consistency enhances our confidence in the anatomical delineation of these subfields within MRI space or the interpretation of ultra-high field functional imaging studies on the hippocampus. Finally, the findings of this study may aid the ongoing efforts of the Hippocampal Subfields Group to create a harmonized hippocampal segmentation protocol (Daugherty et al, 2025; Wisse et al, 2017; Olsen et al, 2019; Yushkevich et al, 2015). Even less conclusive findings, such as the prominent variability in the SUB-CA1 border and the diffuse associations with demographic factors, prompt the need for areas of known variability to be emphasized in training materials. It also highlights the importance of demographic stratifications within datasets, in both data acquisition and analysis.

In the realm of hippocampus research, high-resolution structural MRI provides unparalleled detail to this important region but relies on valid and reliable segmentation. The observations from this study will hopefully contribute to a more data-driven assessment of subfield border locations, including a more diverse and generalizable set of postmortem cases, for hippocampal subfield segmentation protocol development. As the 2024 and the 2019 dataset are publicly available on OpenNeuro and NITRC, respectively, the scientific community is invited to continue the characterization of hippocampal subfield anatomy.

## Footnotes

1. Note, in the current investigation, prosubiculum (located between the subiculum and CA1) is incorporated within the CA1 region-of-interest here and is therefore not further investigated in this section.

## Acknowledgements

We respectfully acknowledge our late coauthors, Ricardo Insausti, Murray Grossman, and John Q. Trojanowski, whose contributions were essential to not only this paper, but the Hippocampal Subfields Group as a whole.

This work was supported by the National Institute of Health grant R01-AG070592, the Swedish Alzheimer Foundation (AF-980872), the Bente Rexed Gerstedt Foundation, the Crafoord Foundation (20230790), and the Biotechnology and Biological Sciences Research Council (BBSRC) (BB/V010549/1). LEMW was supported by MultiPark, a strategic research area at Lund University. The University of Pennsylvania Center for Neurodegenerative Disease Research (CNDR) brain bank is supported by the National Institute on Aging (P30AG072979, P01AG066597, P01AG084497, RF1AG056014).

## References

Adler, D. H., Pluta, J., Kadivar, S., Craige, C., Gee, J. C., Avants, B. B., & Yushkevich, P. A. (2014). Histology-derived volumetric annotation of the human hippocampal subfields in postmortem MRI. Neuroimage, 84, 505–523.

Adler, D. H., Wisse, L. E. M., Ittyerah, R., Pluta, J. B., Ding, S., Xie, L., Wang, J., Kadivar, S., Robinson, J. L., Schuck, T., Trojanowski, J. Q., Grossman, M., Detre, J. A., Elliott, M. A., Toledo, J. B., Liu, W., Pickup, S., Miller, M. I., Das, S. R., Wolk, D. A., & Yushkevich, P. A. (2018). Characterizing the human hippocampus in aging and Alzheimer’s disease using a computational atlas derived from ex vivo MRI and histology. Proceedings of the National Academy of Sciences, 115(16), 4252–4257.

Aggleton, J. P., & Christiansen, K. (2015). The subiculum: The heart of the extended hippocampal system. In Progress in Brain Research (1st ed., Vol. 219). Elsevier B.V. 10.1016/bs.pbr.2015.03.003

Amaral, D. G., & Insausti, R. (1990). Hippocampal formation. In The Human Nervous System (pp. 711–755). Academic Press.

Barbas, H., & Blatt, G. J. (1995). Topographically specific hippocampal projections target functionally distinct prefrontal areas in the rhesus monkey. Hippocampus, 5(6), 511–533. 10.1002/hipo.450050604

Berger, B., Alvarez, C., & Pelaprat, D. (1997). Retrosplenial/presubicular continuum in primates: A developmental approach in fetal macaques using neurotensin and parvalbumin as markers. Developmental Brain Research, 101(1–2), 207–224. 10.1016/S0165-3806(97)00067-9

Berron, D., Vieweg, P., Hochkeppler, A., Pluta, J. B., Ding, S. L., Maass, A., Luther, A., Xie, L., Das, S. R., Wolk, D. A., Wolbers, T., Yushkevich, P. A., Düzel, E., & Wisse, L. E. M. (2017). A protocol for manual segmentation of medial temporal lobe subregions in 7 Tesla MRI. NeuroImage: Clinical, 15(May), 466–482. 10.1016/j.nicl.2017.05.022

Berron, D., Schütze, H., Maass, A., Cardenas-Blanco, A., Kuijf, H. J., Kumaran, D., & Düzel, E. (2016). Strong evidence for pattern separation in human dentate gyrus. Journal of Neuroscience, 36(29), 7569–7579. 10.1523/JNEUROSCI.0518-16.2016

Busch, J. R., Lundemose, S. B., Lynnerup, N., Jacobsen, C., Jørgensen, M. B., & Banner, J. (2019). Post-mortem MRI-based volumetry of the hippocampus in forensic cases of decedents with severe mental illness. Forensic Science, Medicine, and Pathology, 15(2), 213–217. 10.1007/s12024-019-00101-w

Canada, K., Mazloum-Farzaghi, N., Rådman, G., Adams, J. N., Bakker, A., Baumeister, H., Berron, D., Bocchetta, M., Carr, V., Dalton, M. A., de Flores, R., Keresztes, A., La Joie, R., Mueller, S. G., Raz, N., Santini, T., Shaw, T., Stark, C. E. L., Tran, T. T., Wang, L., Wisse, L.E.M., Wuestefeld, A., Yushkevich, P.A., Olsen, R.K., Daugherty, A., on behalf of the Hippocampal Subfields Group (2024). A (sub) field guide to quality control in hippocampal subfield segmentation on high-resolution T 2 -weighted MRI. Human Brain Mapping, 15(19), 1–19. 10.1002/hbm.70004

Chang, W. T., Langella, S. K., Tang, Y., Ahmad, S., Zhang, H., Yap, P. T., Giovanello, K. S., & Lin, W. (2021). Brainwide functional networks associated with anatomically — and functionally — defined hippocampal subfields using ultrahigh-resolution fMRI. Scientific Reports, 11(1), 1–13. 10.1038/s41598-021-90364-7

Dalton, M. A., D’souza, A., Lv, J., & Calamante, F. (2022). New insights into anatomical connectivity along the anterior–posterior axis of the human hippocampus using in vivo quantitative fibre tracking. eLife, 11, 1–29. 10.7554/ELIFE.76143

Dalton, M. A., & Maguire, E. A. (2017). The pre/parasubiculum: a hippocampal hub for scene-based cognition? Current Opinion in Behavioral Sciences, 17, 34–40. 10.1016/j.cobeha.2017.06.001

Dalton, M. A., Zeidman, P., Barry, D. N., Williams, E., & Maguire, E. A. (2017). Segmenting subregions of the human hippocampus on structural magnetic resonance image scans: An illustrated tutorial. Brain and Neuroscience Advances, 1, 239821281770144. 10.1177/2398212817701448

Daugherty, A. M., Carr, V., Canada, K. L., Rådman, G., Brown, T., Augustinack, J., Amunts, K., Bakker, A., Berron, D., Burggren, A., Chetelat, G., de Flores, R., Ding, S.-L., Huang, Y., Insausti, R., Johnson, E., Kanel, P., Keresztes, A., Kedo, O., Kennedy, K. M., Lee, J., Malykhin, N., Martinez, A., Mueller, S., Mulligan, E., Ofen, N., Palombo, D., Pasquini, L., Pluta, J., Raz, N., Riggins, T., Rodrigue, K. M., Saifullah, S., Schlichting, M. L., Stark, C., Wang, L., Yuschkevich, P. A., La Joie, R., Wisse, L. E. M., Olsen, R., on behalf of the Alzheimer’s Disease Neuroimaging Initiative (2025). Harmonized Protocol for Subfield Segmentation in the Hippocampal Body on High-Resolution in vivo MRI from the Hippocampal Subfields Group (HSG). BioRxiv, 2025.04.29.651039. https://www.biorxiv.org/content/10.1101/2025.04.29.651039v1%0Ahttps://www.biorxiv.org/content/10.1101/2025.04.29.651039v1.abstract

de Flores, R., Berron, D., Ding, S., Ittyerah, R., Pluta, J. B., Xie, L., Adler, D. H., Robinson, J. L., Schuck, T., Trojanowski, J. Q., Grossman, M., Liu, W., Pickup, S., Das, S. R., Wolk, D. A., Yushkevich, P. A., & Wisse, L. E. M. (2019). Characterization of hippocampal subfields using ex vivo MRI and histology data: Lessons for in vivo segmentation. Hippocampus, 30(6), 545–564. 10.1002/hipo.23172

de Flores, R., La Joie, R., & Chételat, G. (2015). Structural imaging of hippocampal subfields in healthy aging and Alzheimer’s disease. Neuroscience, 309, 29–50. 10.1016/j.neuroscience.2015.08.033

Dimsdale-Zucker, H. R., Ritchey, M., Ekstrom, A. D., Yonelinas, A. P., & Ranganath, C. (2018). CA1 and CA3 differentially support spontaneous retrieval of episodic contexts within human hippocampal subfields. Nature Communications, 9(1). 10.1038/s41467-017-02752-1

Ding, S-L. (2013). Comparative anatomy of the prosubiculum, subiculum, presubiculum, postsubiculum, and parasubiculum in human, monkey, and rodent. Journal of Comparative Neurology, 521(18), 4145–4162. 10.1002/cne.23416

Ding, S-L., & Van Hoesen, G. W. (2015). Organization and detailed parcellation of human hippocampal head and body regions based on a combined analysis of cyto-and chemoarchitecture. Journal of Comparative Neurology, 523(15), 2233–2253.

Duvernoy, H. M., Cattin, F., & Risold, P. Y. (2005). The human hippocampus: The functional anatomy, vascularization and serial sections with MRI (3rd ed.). Springer.

Duvernoy, H. M., Cattin, F., & Risold, P. Y. (1998). The human hippocampus: The functional anatomy, vascularization and serial sections with MRI. Springer.

Gertz, S. D., Lindenberg, R., & Piavis, G. M. (1972). Structural variationa in the rostral human hippocampus. The Johns Hopkins Medical Journal, 130(6), 367–376.

Grande, X., Wisse, L., & Berron, D. (2023). Ultra-high field imaging of the human medial temporal lobe. In Ultra-High Field Neuro MRI (1st ed., Vol. 10). Elsevier Inc. 10.1016/b978-0-323-99898-7.00031-6

Harrison, P. J. (2004). The hippocampus in schizophrenia: A review of the neuropathological evidence and its pathophysiological implications. Psychopharmacology, 174(1), 151–162. 10.1007/s00213-003-1761-y

Hickling, A. L., Clark, I. A., Wu, Y. I., & Maguire, E. A. (2024). Automated protocols for delineating human hippocampal subfields from 3 Tesla and 7 Tesla magnetic resonance imaging data. Hippocampus, 34(6), 302–308. 10.1002/hipo.23606

Hodgetts, C. J., Voets, N. L., Thomas, A. G., Clare, S., Lawrence, A. D., & Graham, K. S. (2017). Ultra-high-field fMRI reveals a role for the subiculum in scene perceptual discrimination. Journal of Neuroscience, 37(12), 3150–3159. 10.1523/JNEUROSCI.3225-16.2017

Hyman, B. T., Creighton, P. H., Beach, T. G., Bigio, E. H., Cairns, N. J., Carrillo, M. C., Dickson, D. W., Duyckaerts, C., Frosch, M. P., Masilah, E., Mirra, S. S., Nelson, P. T., Schneider, J. A., Thal, D. R., Thies, B., Trojanowski, J. Q., Vinters, H. V, & Montine, T. J. (2012). National Institute on Aging–Alzheimer’s Association guidelines for the neuropathologic assessment of Alzheimer’s disease. Alzheimer’s & Dementia, 8(1), 1–13. 10.1016/j.jalz.2011.10.007.National

Insausti, R., & Amaral, D. G. (2012). Hippocampal formation. In J. Mai & G. Paxinos (Eds.), The Human Nervous System (3rd ed., pp. 896–942). Elsevier Academic Press.

Insausti, R., Muñoz-López, M., Insausti, A. M., & Artacho-Pérula, E. (2017). The human periallocortex: Layer pattern in presubiculum, parasubiculum and entorhinal cortex. A review. Frontiers in Neuroanatomy, 11(October), 1–10. 10.3389/fnana.2017.00084

Kassambara A (2023). rstatix: Pipe-Friendly Framework for Basic Statistical Tests. R package version 0.7.2.

Kerchner, G. A., Hess, C. P., Hammond-Rosenbluth, K. E., Xu, D., Rabinovici, G. D., Kelley, D. A. C., Vigneron, D. B., Nelson, S. J., & Miller, B. L. (2010). Hippocampal CA1 apical neuropil atrophy in mild Alzheimer disease visualized with 7-T MRI. Neurology, 75(15), 1381–1387. 10.1212/WNL.0b013e3181f736a1

Klüver, H., & Barrera, E. (1953). A method for the combined staining of cells and fibers in the nervous system. Journal of Neuropathology & Experimental Neurology, 12(4), 400–403.

Knierim, J. J., & Neunuebel, J. P. (2016). Tracking the flow of hippocampal computation: Pattern separation, pattern completion, and attractor dynamics. Neurobiology of Learning and Memory, 129, 38–49. 10.1016/j.nlm.2015.10.008

La Joie, R., Perrotin, A., de La Sayette, V., Egret, S., Doeuvre, L., Belliard, S., Eustache, F., Desgranges, B., & Chételat, G. (2013). Hippocampal subfield volumetry in mild cognitive impairment, Alzheimer’s disease and semantic dementia. NeuroImage: Clinical, 3, 155–162. 10.1016/j.nicl.2013.08.007

Ledergerber, D., Battistin, C., Blackstad, J. S., Gardner, R. J., Witter, M. P., Moser, M. B., Roudi, Y., & Moser, E. I. (2021). Task-dependent mixed selectivity in the subiculum. Cell Reports, 35(8), 109175.

Leutgeb, J. K, Leutgeb, S., Moser, M. B., & Moser, E. I. (2007). Pattern separation in the dentate gyrus and CA3 of the hippocampus. Science, 315(5814), 961–966.

Mai, J. K., Majtanik, M., & Paxinos, G. (2015). Atlas of the Human Brain. Academic Press.

Marr, D. (1971). Simple memory: a theory for archicortex. Philosophical Transactions of the Royal Society B: Biological Sciences, 262, 23–81.

Maruszak, A., & Thuret, S. (2014). Why looking at the whole hippocampus is not enough-a critical role for anteroposterior axis, subfield and activation analyses to enhance predictive value of hippocampal changes for Alzheimer’s disease diagnosis. Frontiers in Cellular Neuroscience, 8, 1–11. 10.3389/fncel.2014.00095

Moore, M., Hu, Y., Woo, S., O’Hearn, D., Iordan, A. D., Dolcos, S., & Dolcos, F. (2014). A comprehensive protocol for manual segmentation of the medial temporal lobe structures. Journal of Visualized Experiments, 89, 1–8. 10.3791/50991

Moscovitch, M., Cabeza, R., Winocur, G., & Nadel, L. (2016). Episodic memory and beyond: The hippocampus and neocortex in transformation. Annual Review of Psychology, 67, 105–134. 10.1177/0022146515594631.Marriage

Moser, M. B., & Moser, E. I. (1998). Functional differentiation in the hippocampus. Hippocampus, 8(7), 608–619.

Neunuebel, J. P., & Knierim, J. J. (2014). CA3 retrieves coherent representations from degraded input: Direct evidence for CA3 pattern completion and dentate gyrus pattern separation. Neuron, 81(2), 416–427. 10.1016/j.neuron.2013.11.017

Nuninga, J. O., Mandl, R. C. W., Boks, M. P., Bakker, S., Somers, M., Heringa, S. M., Nieuwdorp, W., Hoogduin, H., Kahn, R. S., Luijten, P., & Sommer, I. E. C. (2020). Volume increase in the dentate gyrus after electroconvulsive therapy in depressed patients as measured with 7T. Molecular Psychiatry, 25(7), 1559–1568. 10.1038/s41380-019-0392-6

O’Mara, S. (2005). The subiculum: What it does, what it might do, and what neuroanatomy has yet to tell us. Journal of Anatomy, 207(3), 271–282. 10.1111/j.1469-7580.2005.00446.x

O’Mara, S. M., Commins, S., Anderson, M., & Gigg, J. (2001). The subiculum: A review of form, physiology and function. Progress in Neurobiology, 64(2), 129–155. 10.1016/S0301-0082(00)00054-X

Olsen, R. K., Carr, V. A., Daugherty, A. M., La Joie, R., Amaral, R. S. C., Amunts, K., Augustinack, J. C., Bakker, A., Bender, A. R., Berron, D., Boccardi, M., Bocchetta, M., Burggren, A. C., Chakravarty, M. M., Chételat, G., de Flores, R., DeKraker, J., Ding, S. L., Geerlings, M. I., Huang, Y., Insaustiy, R., Johnson, E. G., Kanel, P., Kedo, O., Kennedy, K. M., Keresztes, A., Lee, J. K., Lindenberger, U., Mueller, S. G., Mulligan, E. M., Ofen, N., Palombo, D. J., Pasquini, L., Pluta, J., Raz, N., Rodrigue, K. M., Schlichting, M. L., Lee Shing, Y., Stark, C. E. L., Steve, T. A., Suthana, N. A., Wang, L., Werkle-Bergner, M., Yushkevich, P. A., Yu, O., & Wisse, L. E. M. (2019). Progress update from the hippocampal subfields group. Alzheimer’s and Dementia: Diagnosis, Assessment and Disease Monitoring, 11, 439–449. 10.1016/j.dadm.2019.04.001.

Poppenk, J., Evensmoen, H. R., Moscovitch, M., & Nadel, L. (2013). Long-axis specialization of the human hippocampus. Trends in Cognitive Sciences, 17(5), 230–240. 10.1016/j.tics.2013.03.005

R Core Team (2025). R: A Language and Environment for Statistical Computing. R Foundation for Statistical Computing, Vienna, Austria. https://www.R-project.org/.

Ravikumar, S., Wisse, L. E. M., Lim, S., Ittyerah, R., Xie, L., Bedard, M. L., Das, S. R., Lee, E. B., Tisdall, M. D., Prabhakaran, K., Lane, J., Detre, J. A., Mizsei, G., Trojanowski, J. Q., Robinson, J. L., Schuck, T., Grossman, M., Artacho-Pérula, E., de Onzoño Martin, M. M. I., del Mar Arroyo Jiménez, M., Muñoz, M., Romero, F. J. M., del Pilar Marcos Rabal, M., Cebada Sánchez, S., Delgado González, J. C., de la Rosa Prieto, C., Parada, M. C., Irwin, D. J., Wolk, D. A., Insausti, R., & Yushkevich, P. A. (2021). Ex vivo MRI atlas of the human medial temporal lobe: characterizing neurodegeneration due to tau pathology. Acta Neuropathologica Communications, 9(1), 1–14. 10.1186/s40478-021-01275-7

Read, M. L., Berry, S. C., Graham, K. S., Voets, N. L., Zhang, J., Aggleton, J. P., Lawrence, A. D., & Hodgetts, C. J. (2024). Scene-selectivity in CA1/subicular complex: Multivoxel pattern analysis at 7T. Neuropsychologia, 194(September 2023), 108783. 10.1016/j.neuropsychologia.2023.108783

Roeske, M. J., Konradi, C., Heckers, S., & Lewis, A. S. (2021). Hippocampal volume and hippocampal neuron density, number and size in schizophrenia: a systematic review and meta-analysis of postmortem studies. Molecular Psychiatry, 26(7), 3524–3535. 10.1038/s41380-020-0853-y

Rosene, D. L., & Van Hoesen, G. W. (1987). The hippocampal formation of the primate brain: a review of some comparative aspects of cytoarchitecture and connections. In E. G. Jones & A. Peters (Eds.), Cerebral cortex (pp. 345–456). Plenum.

Schobel, S. A., Lewandowski, N. M., Corcoran, C. M., Moore, H., Brown, T., Malaspina, D., & Small, S. A. (2009). Differential Targeting of the CA1 Subfield of the Hippocampal Formation by Schizophrenia and Related Psychotic Disorders. Archives of General Psychiatry, 66(9), 938–946. 10.1001/archgenpsychiatry.2009.115

Small, S. A., Schobel, S. A., Buxton, R. B., Witter, M. P., & Barnes, C. A. (2012). A pathophysiological framework of hippocampal dysfunction in ageing and disease. Nature Reviews Neuroscience, 12(10), 585–601. 10.1038/nrn3085.A

Steve, T. A., Yasuda, C. L., Coras, R., Lail, M., Blumcke, I., Livy, D. J., Malykhin, N., & Gross, D. W. (2017). Development of a histologically validated segmentation protocol for the hippocampal body. NeuroImage, 157(June), 219–232. 10.1016/j.neuroimage.2017.06.008

Thom, M. (2014). Review: Hippocampal sclerosis in epilepsy: A neuropathology review. Neuropathology and Applied Neurobiology, 40(5), 520–543. 10.1111/nan.12150

Twait, E. L., Blom, K., Koek, H. L., Zwartbol, M. H. T., Ghaznawi, R., Hendrikse, J., Gerritsen, L., & Geerlings, M. I. (2023). Psychosocial factors and hippocampal subfields: The Medea-7T study. Human Brain Mapping, 44(5), 1964–1984. 10.1002/hbm.26185

Vos De Wael, R., Larivière, S., Caldairou, B., Hong, S. J., Margulies, D. S., Jefferies, E., Bernasconi, A., Smallwood, J., Bernasconi, N., & Bernhardt, B. C. (2018). Anatomical and microstructural determinants of hippocampal subfield functional connectome embedding. Proceedings of the National Academy of Sciences of the United States of America, 115(40), 10154–10159. 10.1073/pnas.1803667115

Wisse, L. E. M., Biessels, G. J., Stegenga, B. T., Kooistra, M., van der Veen, P. H., Zwanenburg, J. J. M., van der Graaf, Y., & Geerlings, M. I. (2015). Major depressive episodes over the course of 7 years and hippocampal subfield volumes at 7 tesla MRI: The PREDICT-MR study. Journal of Affective Disorders, 175, 1–7.

Wisse, L. E. M., Daugherty, A. M., Olsen, R. K., Berron, D., Carr, V. A., Stark, C. E. L., Amaral, R. S. C., Amunts, K., Augustinack, J. C., Bender, A. R., Bernstein, J. D., Boccardi, M., Bocchetta, M., Burggren, A., Chakravarty, M. M., Chupin, M., Ekstrom, A., de Flores, R., Insausti, R., … la Joie, R. (2017). A harmonized segmentation protocol for hippocampal and parahippocampal subregions: Why do we need one and what are the key goals? Hippocampus, 27(1), 3–11. 10.1002/hipo.22671

Wuestefeld, A., Baumeister, H., Adams, J. N., de Flores, R., Hodgetts, C. J., Mazloum-Farzaghi, N., Olsen, R. K., Puliyadi, V., Tran, T. T., Bakker, A., Canada, K. L., Dalton, M. A., Daugherty, A. M., La Joie, R., Wang, L., Bedard, M. L., Buendia, E., Chung, E., Denning, A., … Wisse, L. E. M. (2024). Comparison of histological delineations of medial temporal lobe cortices by four independent neuroanatomy laboratories. Hippocampus, 34(5), 241–260. 10.1002/hipo.23602

Yushkevich, P. A., Muñoz López, M., Iñiguez De Onzoño Martin, M. M., Ittyerah, R., Lim, S., Ravikumar, S., Bedard, M. L., Pickup, S., Liu, W., Wang, J., Hung, L. Y., Lasserve, J., Vergnet, N., Xie, L., Dong, M., Cui, S., McCollum, L., Robinson, J. L., Schuck, T., de Flores, R., Grossman, M., Tisdall, M. D., Prabhakaran, K., Mizsei, G., Das, S. R., Artacho-Péula, E., Del Mar Arroyo Jiménez, M., Marcos Raba, M. P., Molina Romero, F. J., Cebada Sánchez, S., Delgado González, J. C., De La Rosa-Pietro, C., Córcoles Parada, M., Lee, E. B., Trojanowski, J. Q., Ohm, D. T., Wisse, L. E. M., Wolk, D. A., Irwin, D. J., & Insausti, R. (2021). Three-dimensional mapping of neurofibrillary tangle burden in the human medial temporal lobe. Brain, 144(9), 2784–2797. 10.1093/brain/awab262

Yushkevich, P. A., Amaral, R. S. C., Augustinack, J. C., Bender, A. R., Bernstein, J. D., Boccardi, M., Bocchetta, M., Burggren, A. C., Carr, V. A., Chakravarty, M. M., Chételat, G., Daugherty, A. M., Davachi, L., Ding, S. L., Ekstrom, A., Geerlings, M. I., Hassan, A., Huang, Y., Iglesias, J. E., Kerchner, G. A., La Rocque, K. F., Libby, L. A., Malykhin, N., Mueller, S. G., Olsen, R. K., Palombo, D. J., Parekh, M. B., Pluta, J. B., Preston, A. R., Pruessner, J. C., Ranganath, C., Raz, N., Schlichting, M. L., Schoemaker, D., Singh, S., Stark, C. E. L., Suthana, N., Tompary, A., Turowski, M. M., Van Leemput, K., Wagner, A. D., Wang, L., Winterburn, J. L., Wisse, L. E. M., Yassa, M. A., & Zeineh, M. M. (2015). Quantitative comparison of 21 protocols for labeling hippocampal subfields and parahippocampal subregions in in vivo MRI: Towards a harmonized segmentation protocol. NeuroImage, 111, 526–541. 10.1016/j.neuroimage.2015.01.004

Yushkevich, P. A., Piven, J., Hazlett, H. C., Smith, R. G., Ho, S., Gee, J. C., & Gerig, G. (2006). User-guided 3D active contour segmentation of anatomical structures: Significantly improved efficiency and reliability. NeuroImage, 31(3), 1116–1128. 10.1016/j.neuroimage.2006.01.015

Zeidman, P., & Maguire, E. A. (2016). Anterior hippocampus: The anatomy of perception, imagination and episodic memory. Nature Reviews Neuroscience, 17(3), 173–182. 10.1038/nrn.2015.24

Zeineh, M. M., Chen, Y., Kitzler, H. H., Hammond, R., Vogel, H., & Rutt, B. K. (2015). Activated iron-containing microglia in the human hippocampus identified by magnetic resonance imaging in Alzheimer disease. Neurobiology of Aging, 36(9), 2483–2500. 10.1016/j.neurobiolaging.2015.05.022

Zeineh, M. M., Engel, S. A., & Bookheimer, S. Y. (2000). Application of cortical unfolding techniques to functional MRI of the human hippocampal region. NeuroImage, *11*(6 I), 668–683. 10.1006/nimg.2000.0561

Zeineh, M. M., Engel, S. A., Thompson, P. M., & Bookheimer, S. Y. (2001). Unfolding the human hippocampus with high resolution structural and functional MRI. The Anatomical Record, 265(2), 111–120. 10.1002/ar.1061

