## Supplementary material for "Characterization of hippocampal subfields using histology-based annotated postmortem MRI: Lessons for in vivo segmentation II"

**SUPPLEMENTAL MATERIAL**

**Supplementary methods** 2

**Supplementary methods: Section 3.1. The order of appearance of hippocampal subfields along the long axis of the hippocampal head** 2

**Supplementary methods: Section 3.2. The order of disappearance of hippocampal subfields along the long axis of the hippocampal tail** 2

**Supplementary methods: Section 3.3. The order of appearance and disappearance of subicular complex subregions in the hippocampus** 3

**Supplementary methods & results: Section 3.8. The order of appearance and disappearance of subicular complex subregions in the hippocampus** 3

**Supplementary figures** 4

**Supplementary Figure 1.** 4

**Supplementary Figure 2.** 5

**Supplementary Figure 3.** 6

**Supplementary Figure 4.** 7

**Supplementary Figure 5.** 8

**Supplementary Figure 6.** 9

**Supplementary tables** 10

**Supplementary Table 1.** 10

**Supplementary Table 2.** 11

**Supplementary Table 3.** 12

**Supplementary Table 4.** 13

**Supplementary Table 5.** 14

**Supplementary Table 6.** 15

**Supplementary Table 7**. 16

### Supplementary methods

#### Supplementary methods: Section 3.1. The order of appearance of hippocampal subfields along the long axis of the hippocampal head

In a number of specimens, segmentation inconsistencies were observed. These were particularly noticeable for the CA2 subfield, where segmentations appeared, then disappeared (being replaced with the segmentation of another subfield), and subsequently reappeared in five specimens—four of which were included in this analysis (CNDR13, CNDR15, CNDR22, and HNL04). Such inconsistencies were primarily due to minor inaccuracies in the registration process used to align segmentations within the full MRI space, rather than inaccuracies in the histological annotations. For the specimens affected, since the segmentations did not actually disappear, the first appearance of CA2 was regarded as its starting point. In specimen CNDR13, however, the first appearance of CA2 was deemed incorrect by R. I., with the subsequent reappearance considered its true appearance slice.

With regard to specimen inclusion and exclusion, CNDR10 was included by De Flores et al. (2019) but excluded from the current study due to uncertainty about the exact starting point of CA3. A final note regards to the exceptions to the 'five-slices rule' mentioned in the approach. Although there were five or more consecutive slices with insufficient coverage anterior to the appearance of CA3 in CNDR24 and CA2 in HNL05, their appearance could be inferred based on neuroanatomical knowledge. These cases were therefore included in the study.

#### Supplementary methods: Section 3.2. The order of disappearance of hippocampal subfields along the long axis of the hippocampal tail

In two cases (CNDR10 and CNDR21), the DG was visible on MRI but not segmented 2 mm anterior to the most posterior slice. Hence, we defined the DG based on MRI features. For CNDR21, the SUB was also not segmented 2 mm prior to the most posterior slice, but its presence could be inferred from the adjacent slices. Additionally, in one specimen (CNDR19), the CA1 subfield was not segmented 1 mm anterior to the most posterior slice; nevertheless, its presence could be inferred from the preceding slices. With regard to specimen inclusion and exclusion, CNDR02 was excluded from the analysis focusing on the most posterior 1 mm of the tail because tissue to be analysed was partially cut off in this slice. Note that this case was included by De Flores et al. (2019).

**Supplementary methods: Section 3.3. The order of appearance and disappearance of subicular complex subregions in the hippocampus**

For one case (CNDR22), some histological sections were partially missing in the hippocampal body, which meant that the precise appearance point of the parasubiculum could not be determined. As there were only 4 slices (< 1mm) between the start of these missing sections and the appearance of parasubiculum this case was kept in the analysis.

**Supplementary methods & results: Section 3.8. The order of appearance and disappearance of subicular complex subregions in the hippocampus**

*Approach:* For certain cases, the SUB-CA1 border was more medial than the medial anchor point, in which case the relative SUB-CA1 border was reported as a negative percentage (see Supplemental Figure 6). Moreover, the approach was adjusted slightly for cases which were rotated. A line was drawn between the two voxels most distal from each other in the hippocampal segmentation; this line was used as the horizontal axis to draw the rest of the referential lines.

### Supplementary figures

**Supplementary Figure 1.** Comparison of coronal sections of the hippocampus, sliced perpendicular to the long axis of the hippocampus (upper row) and when the hippocampus is rotated at a 25° angle. The data displayed here are the template and average segmentation of the post-mortem hippocampus atlas in Adler et al. (2018). Displayed sections are taken at the same location for both angulations, starting 4 mm after the most anterior tip of the hippocampus, with 4 mm spacing between consecutive sections.

As can be observed in figures a-j, the slicing angulation affects the appearance of the hippocampus in several ways as indicated by the white arrows. In the first section, the observed difference is in the medial portion where in figure f, in contrast to figure a, the hippocampus includes the medial portion of the cortex. In figure g subfield cornu ammonis (CA) 3 is present in addition to CA2, whereas only CA2 is present in figure b. In figure h the medial portion of the hippocampus already starts to show a body-like appearance with an x-shaped appearance of CA3, which is not the case in c. Figure d is still in the head of the hippocampus indicated by the presence of the uncal apex whereas i already represents a body slice. In e and j the coronal sections both represent body sections with a similar composition of the subfields. This indicates that the slicing angle in atlases could potentially affect features that are often used for the development of subfield protocols, such as the composition and location of subfields in coronal sections but also the appearance of subfields in relation to the longitudinal axis of the hippocampus in metrics such as millimeters.

It should be noted that this is not a quantitative comparison and this comparison is therefore not definitive. The composition of subfields might look more similar if different sections were compared rather than a set distance from the anterior tip of the hippocampus. CA = cornu ammonis, DG = dentate gyrus, SRLM = stratum radiatum lacunosum moleculare, SUB = subiculum

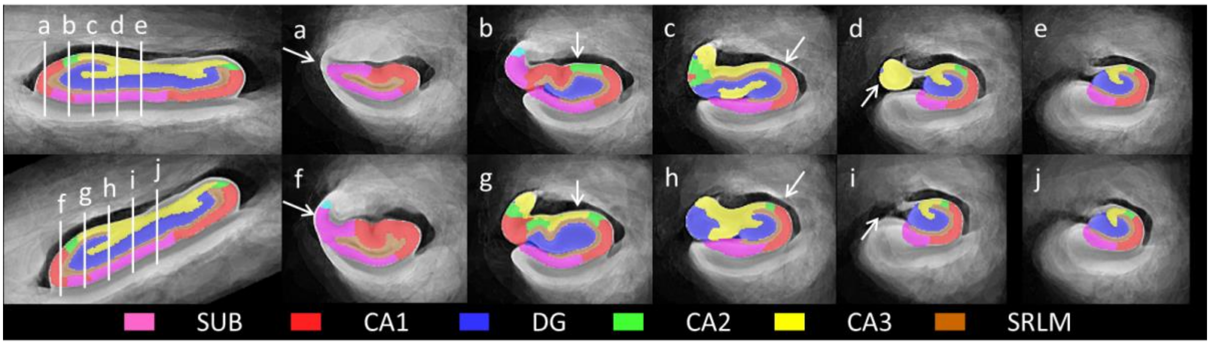

**Supplementary Figure 2.** An example of a specimen (NDRI14) that was excluded due to uncertainty regarding the exact starting point of a subfield, in this case, CA3. Images *a–e* displays five consecutive slices in an anterior-posterior direction, leading up to the appearance of CA3 in f. The slices preceding the first labelled CA3 slice did only show partial coverage of the segmentation, which potentially obscured the identification of a more anterior emergence of CA3. Note that CA2 had already appeared prior to image *a*, so its emergence was recorded earlier. CA = cornu ammonis, DG = dentate gyrus, SUB = subiculum.

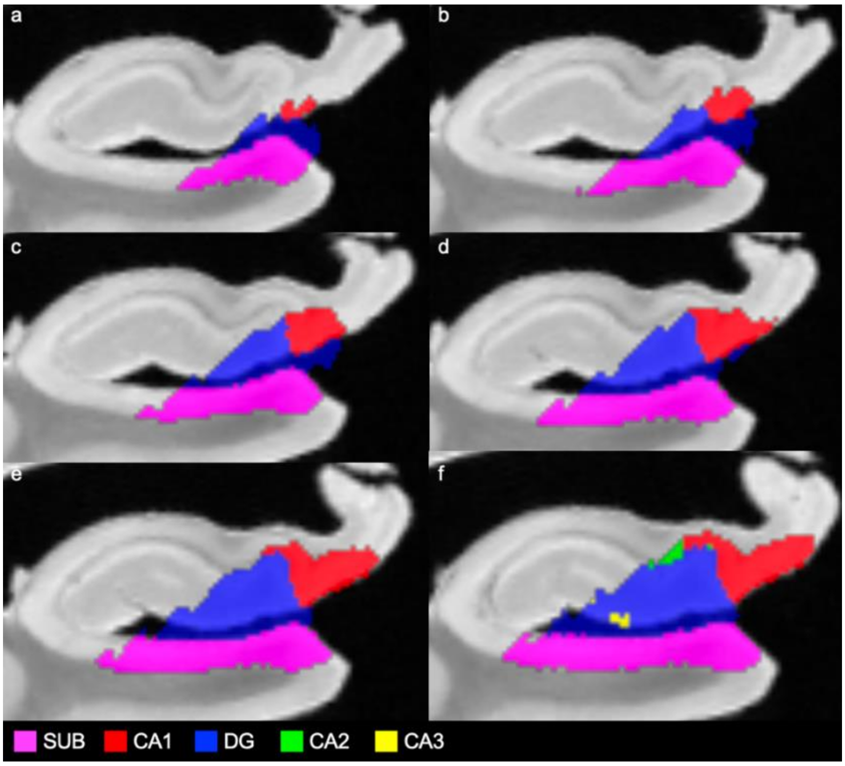

**Supplementary Figure 3.** Two examples of the determination of the most posterior slice of the hippocampal tail. In both rows, images 1–5 depict slices in an anterior-to-posterior direction: 3 mm anterior to the most posterior slice, 2 mm anterior, 1 mm anterior, the most posterior slice, and the slice following it, respectively. In row a (HNL04), the tail splits into two distinct portions (pink and red arrows). In this type of tail, the superior portion (pink arrow) was followed to identify the most posterior slice, which was defined as the final slice where grey matter was visible (blue arrow). In row b (CNDR16), the tail is characterized by grey matter 'blobs' separating laterally from the hippocampus (green arrow). Here, the separating 'blobs' were followed to identify the most posterior slice, which was defined as the final slice where tissue separation was observed (yellow arrow).

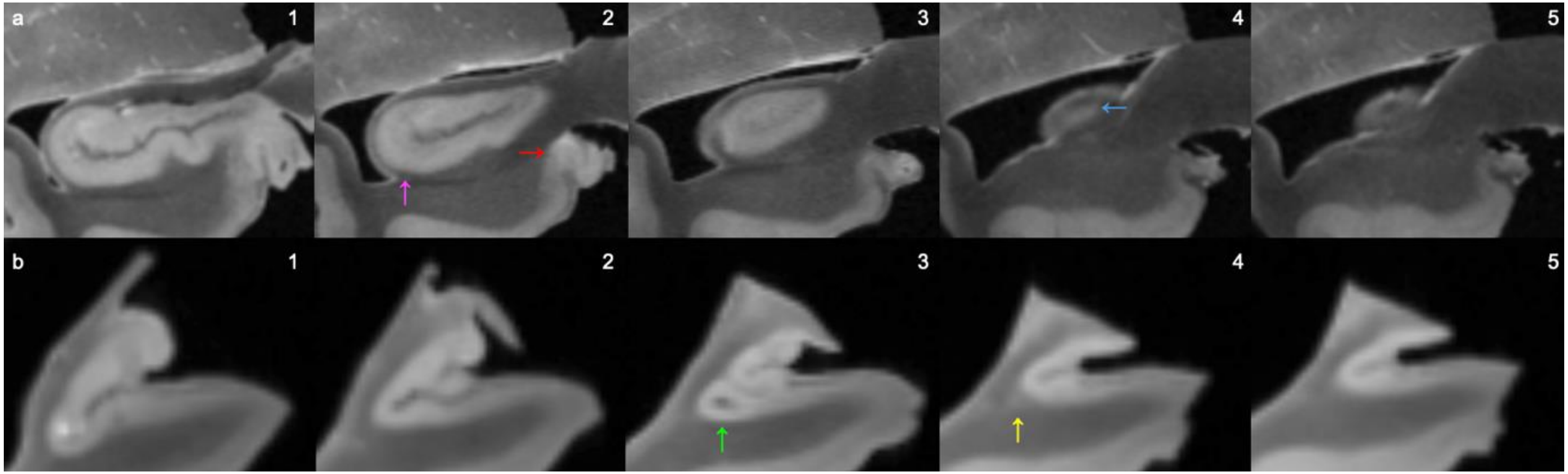

**Supplementary Figure 4.** Volume of CA1, CA2, and CA3 at varying anterior-posterior locations in the hippocampal body. CA = cornu ammonis,

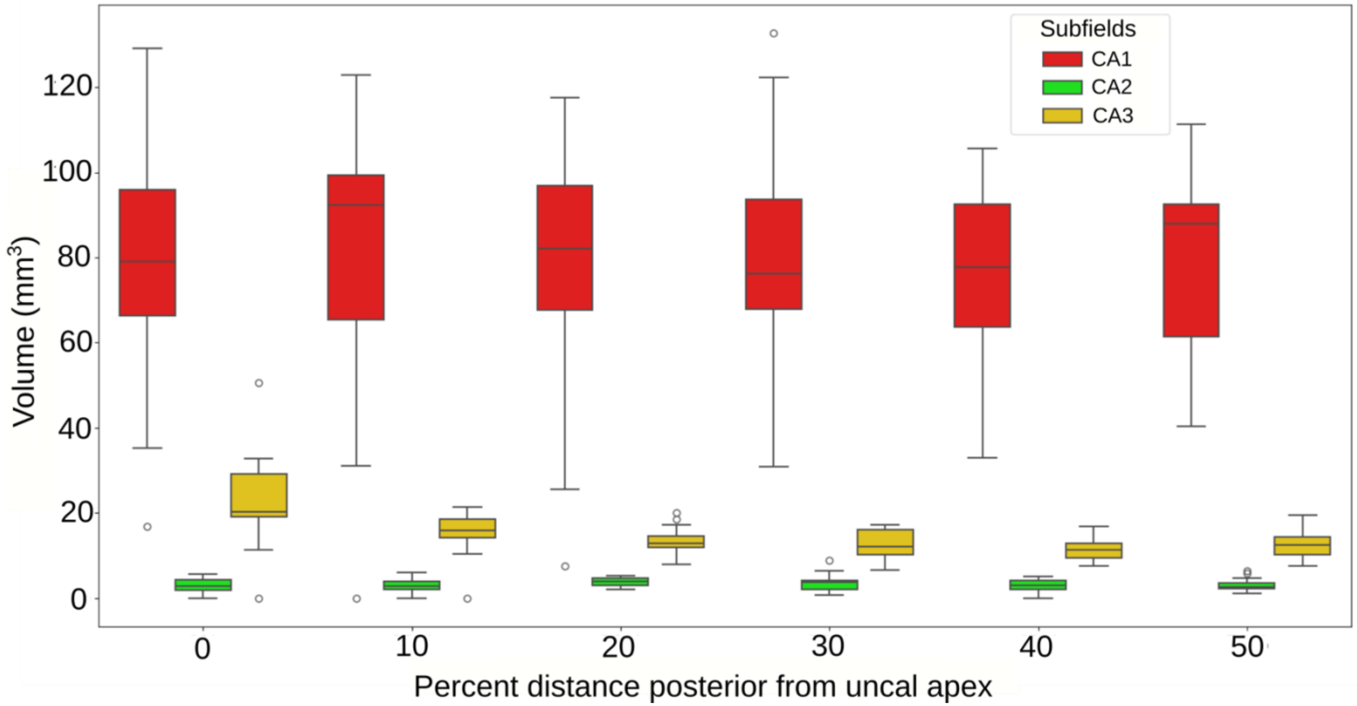

**Supplementary Figure 5.** Scatterplots showing the Spearman's rank correlations between hippocampal length measures (head, total, and ratio) and age and between hippocampal head length and total hippocampal length.

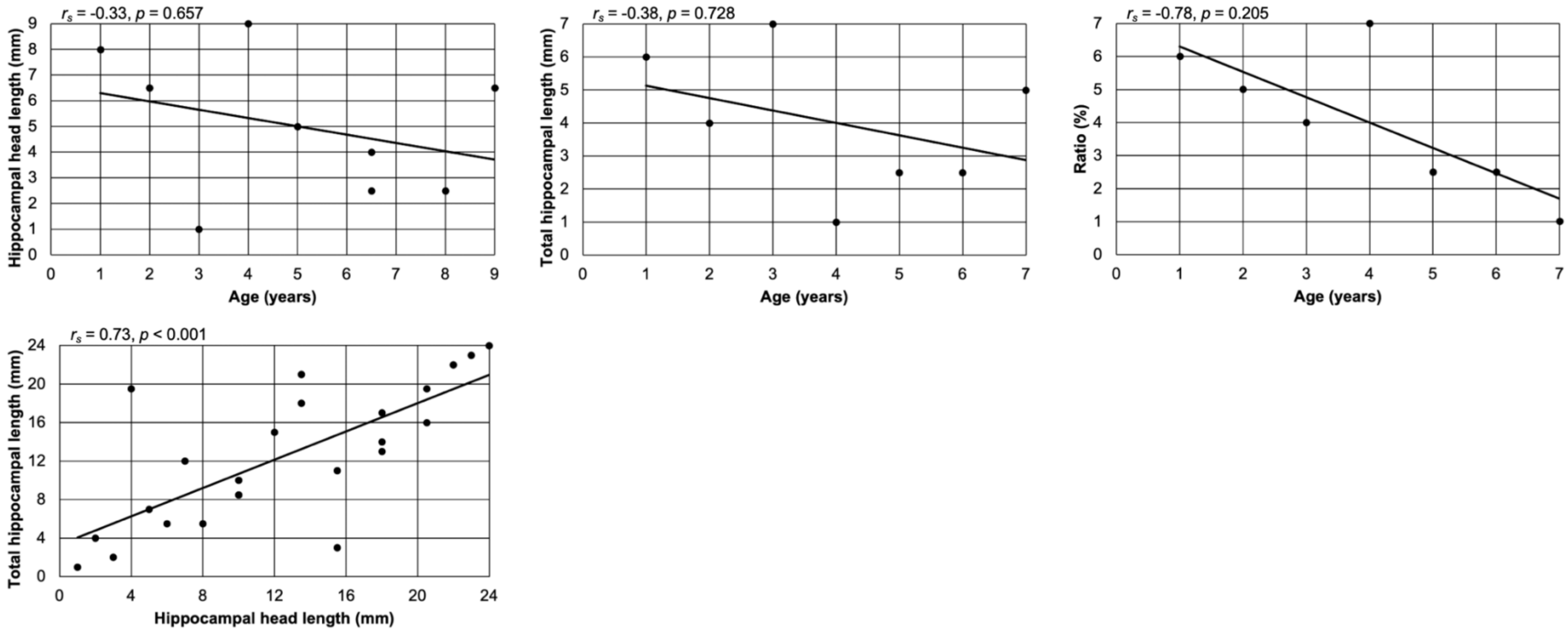


$r_s$ = Spearman's rank correlation coefficient, $p$ = $p$-value

**Supplementary Figure 6.** Example of a SUB-CA1 border more medial than the anchorpoint, thus resulting in a negative percentage measurement. Moreover, this figure exemplifies a case where the most medial point of CA3 is more medial than the most medial point of the DG. In this case, the most medial point of CA3 (white circle) determines the medial anchorpoint (arrow). CA = cornu ammonis, DG = dentate gyrus, SUB = subiculum

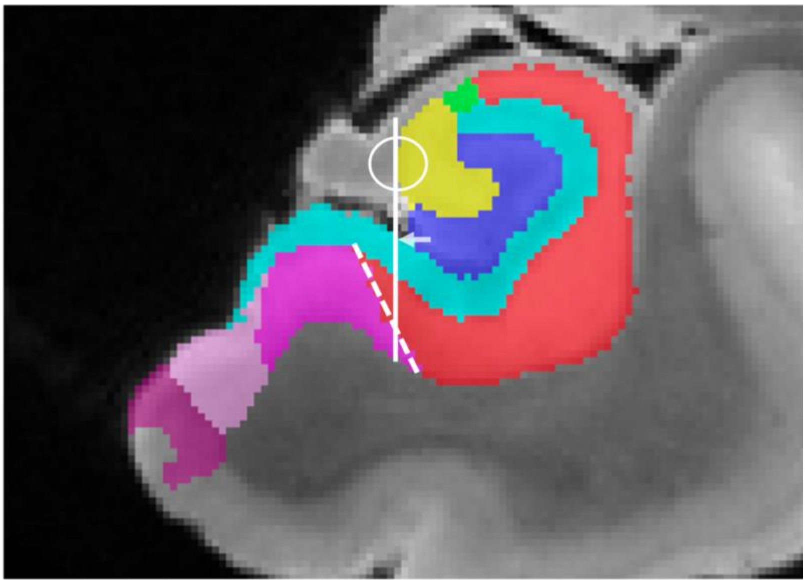

### Supplementary tables

**Supplementary Table 1.** Overview of which dataset and which cases were used for each of the research questions.

| Dataset | Specimen | Section | | | | | | | |
|---|---|---|---|---|---|---|---|---|---|
| | | 3.1 | 3.2 | 3.3 | 3.4 | 3.5 | 3.6 | 3.7 | 3.8 |
| 2019 | NDRI01[a] | | X | | | | | X | X |
| 2019 | NDRI02 | X | | | | | | X | |
| 2019 | NDRI05 | | | | | | | X | X |
| 2019 | NDRI08 | X | X | | | | | X | X |
| 2019 | NDRI09 | | X | | | | | X | X |
| 2019 | NDRI12 | | X | | | | | X | X |
| 2019 | NDRI14 | | | | | | | X | X |
| 2019 | CNDR02 | X | X | | | | | X | X |
| 2019 | CNDR10 | | X | | | | | X | X |
| 2024 | CNDR13 | X | X | X | X | | X | X | X |
| 2024 | CNDR14 | X | X | X | X | X | X | X | X |
| 2024 | CNDR15 | X | X | X | X | | X | X | X |
| 2024 | CNDR16 | X | X | X | X | X | X | X | X |
| 2024 | CNDR17 | X | X | X | X | X | X | X | X |
| 2024 | CNDR18 | | | | | | X | X | X |
| 2024 | CNDR19 | | X | | | X | X | X | |
| 2024 | CNDR20 | X | X | X | X | X | X | X | X |
| 2024 | CNDR21 | X | X | X | X | X | X | X | X |
| 2024 | CNDR22 | X | X | X | X | X | X | X | X |
| 2024 | CNDR23 | X | | X | X | X | X | X | X |
| 2024 | CNDR24 | X | X | X | X | | X | X | X |
| 2024 | HNL01 | X | | X | | | | X | X |
| 2024 | HNL02 | X | | X | X | | X | X | X |
| 2024 | HNL03 | | | | | X | | X | X |
| 2024 | HNL04 | X | | X | X | X | X | X | X |
| 2024 | HNL05 | X | | X | X | x | x | X | X |

**Supplementary Table 2.** The mean ± SD distances between subfields, presented in millimetres (mm) and as a percentage of head length (%), for both atlases. These distances correspond to the standard order of subfield progression observed.

| **2024 atlas** | | | **2019 atlas** | | |
|---|---|---|---|---|---|
| **Distance** | **mm** | **% of head length** | **Distance** | **mm** | **% of head length** |
| **SUB to CA1** | 1.9 ± 0.6 | 11.3 ± 3.2 | SUB to CA1 | 1.7 ± 0.5 | 9.1 ± 3.0 |
| **CA1 to DG** | 3.7 ± 1.7 | 22.1 ± 9.6 | CA1 to DG | 4.4 ± 0.5 | 23.8 ± 4.0 |
| **DG to CA3** | 1.6 ± 1.6 | 9.5 ± 9.7 | DG to CA2 | 0.4 ± 0.3 | 2.1 ± 1.8 |
| **CA3 to CA2** | 3.0 ± 1.7 | 18.8 ± 11.2 | CA2 to CA3 | 2.3 ± 1.9 | 12.0 ± 9.5 |
| **CA2 to uncal apex** | 6.3 ± 2.4 | 38.3 ± 13.8 | CA3 to uncal apex | 9.8 ± 0.7 | 53.0 ± 7.1 |

SUB = subiculum, CA = cornu ammonis, DG = dentate gyrus.

**Supplementary Table 3.** Mean percent of CA1, CA2, and CA3 around the SRLM lateral to the most medial border of the DG.

| Percent distance from posterior uncal apex | CA1 | CA2 | CA3 |
|---|---|---|---|
| 0% (most anterior) | 81.1% | 8.7% | 4.9% |
| 10% | 83.1% | 8.7% | 4.8% |
| 20% | 84.0% | 9.7% | 3.8% |
| 30% | 83.1% | 9.5% | 4.1% |
| 40% | 85.0% | 9.2% | 3.3% |
| 50% (most posterior) | 84.6% | 9.6% | 3.3% |

CA = cornu ammonis, SRLM = stratum radiatum lacunosum moleculare.

**Supplementary Table 4.** Mean percent proportion of CA1, CA2, and CA3 around the curved SRLM length from de Flores et al. (2019) and Steve et al. (2017), and the mean encompassment of the SRLM *volume* for the same subfield.

| **Subfield** | **CA1** | **CA2** | **CA3** |
|---|---|---|---|
| **de Flores et al. (2019)** | 47.0% | 10.6% | 9.4% |
| **Steve et al. (2017)** | 68.7% | 19.1% | 2.5% |
| **Current study (volume proportion of SRLM)** | 83.5% | 9.21% | 4.0% |

CA = cornu ammonis, SRLM=stratum radiatum lacunosum moleculare

**Supplementary Table 5.** Mean ± SD (range) for hippocampal head length, total length, and the head-to-total length ratio in the dementia and non-dementia groups, as well as by sex.

| | **Dementia status** | | | | **Sex** | | | |
|---|---|---|---|---|---|---|---|---|
| | ***n*** | **Dementia** | ***n*** | **Non-dementia** | ***n*** | **Men** | ***n*** | **Women** |
| Head length (mm) | 17 | 16.0 ± 2.1 (11.8–19.8) | 9 | 17.3 ± 1.3 (16.0–20.2) | 5 | 17.8 ± 1.6 (16.0–20.2) | 4 | 16.6 ± 0.5 (16.2–17.2) |
| Total length (mm) | 17 | 41.3 ± 3.5 (34.6–46.0) | 7 | 42.5 ± 4.2 (36.4–49.6) | 4 | 43.7 ± 5.5 (36.4–49.6) | 3 | 40.9 ± 1.2 (40.2–42.2) |
| Ratio (%) | 17 | 38.8 ± 3.3 (32.1–43.1) | 7 | 41.3 ± 2.5 (39.1–46.7) | 4 | 42.0 ± 3.3 (39.1–46.7) | 3 | 40.5 ± 0.3 (40.3–40.8) |

*n* = number of cases.

**Supplementary Table 6.** Statistical comparison between the original rater from De Flores et al (2019) with the current rater on the measurements of the relative position of the SUB-CA1 border along the hippocampal and DG width, measured from medial (0%) to lateral (100%).

| | **2019 dataset – 2019 rater** | | **2019 dataset – current rater** | | ***p*-value** |
|---|---|---|---|---|---|
| **Reference** | **0 mm mean (*SD*)** | **20 mm mean (*SD*)** | **0 mm mean (*SD*)** | **20 mm mean (*SD*)** | |
| **% Of CA+DG width** | 46.5 (14.2) | 29.4 (26.8) | 43.6 (15.1) | 17.6 (18.2) | 0.35 |
| **% of DG width** | 60.7 (17.7) | 32.6 (30) | 56.7 (17.9) | 22.6 (23.4) | 0.26 |

Paired t-test comparison between the 2019 and current rater. CA=cornu ammonis, DG=dentate gyrus, SUB=subiculum.

**Supplementary Table 7**. SUB-CA1 border location and its relationship with dementia status.

| Dataset | Reference | Distance from the uncus | Dementia status | | | | |
|---|---|---|---|---|---|---|---|
| | | | Dementia | | Non-dementia | | *p* |
| | | | *M* | *SD* | *M* | *SD* | |
| **2019** | **CA+DG** | **0mm** | 40.2 | 11.5 | 49.3 | 21.4 | >0.99 |
| | | **20mm** | 12.8 | 18.7 | 25.6 | 17.5 | 0.79 |
| | **DG** | **0mm** | 52 | 12.5 | 64.5 | 25.7 | >0.99 |
| | | **20mm** | 15.8 | 23.7 | 33.8 | 22.2 | 0.79 |
| **2024** | **CA+DG** | **0mm** | 18.8 | 28.3 | 4 | 12.4 | 0.9 |
| | | **20mm** | 5.5 | 14.5 | -35.8 | 54.5 | 0.79 |
| | **DG** | **0mm** | 24.4 | 36.7 | 6 | 17.8 | 0.9 |
| | | **20mm** | 9.1 | 23.4 | -51.5 | 75.1 | 0.79 |
| **2019 + 2024** | **CA+DG** | **0mm** | 25.5 | 26 | 21 | 27.7 | 0.49 |
| | | **20mm** | 7.78 | 15.7 | -12.8 | 52.9 | 0.79 |
| | **DG** | **0mm** | 33.1 | 33.4 | 27.9 | 35.9 | 0.61 |
| | | **20mm** | 11.2 | 22.9 | -19.5 | 72.9 | 0.7 |

*M* = Mean in percent, *SD* = Standard deviation, $r_s$ = Spearman's rank correlation coefficient. CA = cornu ammonis, DG = dentate gyrus. All *p*-values are FDR-corrected.
